# Adenosine-specific transcriptional programs in murine connective tissue type mast cells

**DOI:** 10.1101/2025.11.10.687562

**Authors:** Qihua Liang, Volodymyr Tsvilovskyy, Anouar Belkacemi, Merima Bukva, Christin Richter, Nicole Ludwig, Andreas Keller, Marc Freichel

**Affiliations:** Institute of Pharmacology, Heidelberg University, Heidelberg, Germany; DZHK (German Centre for Cardiovascular Research), partner site Heidelberg/Mannheim, Heidelberg, Germany; Clinical Bioinformatics, Saarland University; Helmholtz Institute for Pharmaceutical Research Saarland, Helmholtz Center for Infection Research, Saarbrücken, Germany; PharmaScienceHub, Saarland University, Saarbrücken, Germany; Department of Human Genetics, Saarland University, Homburg/Saar, Germany

**Keywords:** peritoneal mast cells, adenosine, compound 48/80, antigen, calcium, transcriptome analysis

## Abstract

Mast cells are tissue-resident immune cells that are critical for the pathogenesis of allergic and inflammatory disorders. In addition to classic host defense against parasites, food quality control through antigen avoidance has been recently elucidated as additional physiological mast cell function. The purine nucleoside adenosine (ADO), like other mast cell activators, such as antigens or Mrgprb2 agonists, increases intracellular Ca^2+^ concentration; however, it fails to induce degranulation of preformed mediators when applied to isolated mast cells alone, and there is limited knowledge of whether ADO evokes the de novo synthesis and release of inflammatory mediators such as cytokines in tissue mast cells. An unbiased genome-wide analysis of gene expression triggered by various mast cell activators should enable identification of the gene program specifically activated by adenosine in mast cells and thereby reveal new components of the associated inflammatory responses. To this end, we performed bulk RNA sequencing (RNA-Seq) in primary murine peritoneal mast cells (PMCs) as connective tissue mast cells. By comparing responses evoked by ADO stimulation with those of the Mrgprb2 compound 48/80 and antigens that activate FcεRI receptors, we identified 393 genes whose expression was uniquely regulated by ADO, including upregulation of genes encoding de novo synthesized mediators such as *Tgfa* and *Il7*. Transcription factor activity inference, protein classification, functional enrichment analysis, protein-protein interaction network, and topology analysis revealed that these genes play roles in phosphoinositide signaling, vesicle trafficking, glycolysis, mitochondrial activity, and cell cycle arrest. In summary, our work elucidates a distinct ADO-triggered response and identifies a set of proteins including de novo synthesized mediators whose functional relevance in adenosine-evoked mast cell activation and inflammatory reactions can be evaluated in future studies.

## 1 Introduction

Mast cells are ancient, evolutionarily conserved resident cells in many tissues. They are located at the host’s environmental interfaces, such as the skin, respiratory tract, and gastrointestinal lining, and act as critical sentinels of the immune system.^1,2^ While traditionally viewed for their detrimental roles in allergic disorders and asthma^3,4^, as well as in itch^5^ and food allergy^6^, mast cells have increasingly been recognized for their beneficial functions in wound healing, host defense, and antigen avoidance.^7–10^

A morphologic feature of mast cells is their abundance of electron-dense secretory granules, which contain large amounts of preformed compounds, including biogenic amines (histamine and serotonin)^11,12^, specific preformed cytokines (for example, tumor necrosis factor and vascular endothelial growth factor)^13,14^, serglycin proteoglycans^15^, various lysosomal enzymes^16^, and many mast cell-specific proteases^17,18^. In mice, mast cells are classified into two major subtypes: connective tissue mast cells (CTMCs) and mucosal mast cells (MMCs). Peritoneal mast cells, the focus of our study, correspond to CTMC subtype.^1,19–22^ CTMCs predominantly express two distinct chymases: the β-chymase, mouse mast cell protease 4 (mMCP-4), and the α-chymase, mMCP-5. In addition, they express tryptases mMCP-6 and mMCP-7, as well as carboxypeptidase A3 (CPA3).^22–24^

Mast cell activation follows a tri-phasic cascade. The immediate response, occurring within seconds to minutes, involves the degranulation and release of the above-mentioned preformed compounds into the extracellular space.^25^ The rapid, intermediate-phase synthesis of lipid mediators follows this. To this end, enzymes process membrane phospholipids to generate arachidonic acid derivatives, such as leukotrienes and prostaglandins.^26^ Finally, the late-phase response is initiated as transcription factors, notably NFAT and NF-κB, drive the *de novo* synthesis of a wide array of cytokines and chemokines.^27,28^

Adenosine (ADO) signals to cells in an autocrine or paracrine manner. Mast cells show either anti-inflammatory or pro-inflammatory responses upon ADO stimulation, depending on the receptor engaged. There are four distinct G protein-coupled receptor subtypes for ADO: A1, A2a, A2b, and A3. Of these, A2a and A2b receptors are coupled to G alpha (s) protein. They lead to the activation of adenylyl cyclase (AC) and increase the production of cAMP. In contrast, A1 and A3 receptors inhibit AC and are coupled to G alpha (i), leading to a reduction of cAMP production. Moreover, A2b and A3 can couple to G alpha (q) protein, activating phospholipase C (PLC).^29^ PLC catalyzes the hydrolysis of phosphatidylinositol 4,5-bisphosphate (PIP2) into diacylglycerol (DAG) and inositol 1,4,5-trisphosphate (IP3), with IP3 initiating Ca^2+^ release from the endoplasmic reticulum (ER). As with other Ca^2+^-mobilizing pathways, ER depletion triggers calcium entry. In a mast cell/basophil cell line, ADO receptor agonist NECA was shown to activate a class of Ca^2+^ entry channels named “Calcium Release Activated channels” (CRAC).^30^ Activation of A2a is predominantly reported to be anti-inflammatory, whereas signaling through A1, A2b, and A3 can be either pro- or anti-inflammatory.^29,31^

ADO acts synergistically with canonical stimuli; when combined with antigen challenge, it enhances FcεRI-mediated degranulation. However, ADO alone is insufficient to trigger significant degranulation. ADO has been shown to potentiate mediator release from mast cells upon antigen stimulation.^32–34^ In the human HMC-1 mast cell line, IL-8 release can be evoked by ADO-receptor stimulation; however, data from different research groups are controversial.^35–38^

The classical immunologic pathway for mast cell activation involves the binding of an antigen to IgE antibodies, which are already bound to the mast cell’s high-affinity receptor for IgE (FcεRI). This receptor activation induces downstream signaling cascades mediated by PLC.^39^ The subsequent depletion of intracellular Ca^2+^ activates Ca^2+^ influx through plasma membrane channels that partially depend on Orai1 expression, leading to a robust increase in cytosolic Ca^2+^ that drives degranulation and the *de novo* synthesis of inflammatory mediators.^40^ For example, antigen-evoked release of TNF-alpha and IL-6 was partially reduced in Orai1-deficient primary mast cells.^40^

In addition, mast cells can degranulate via activation of the Mas-related G protein–coupled receptor b2 (*Mrgprb2*), the ortholog of human MRGPRX2.^41,42^ Activation of Mrgprb2 initiates a signaling cascade involving activation of the PLC-IP3 axis, followed by an intracellular Ca^2+^ rise and mast cell degranulation, along with the de novo synthesis of inflammatory mediators.^40^

The pathological significance of ADO signaling is evident in several human inflammatory disorders. Notably, inhaled ADO provokes bronchoconstriction in patients with asthma or chronic obstructive pulmonary disease but not in healthy individuals.^43^ Consistent with a role in cutaneous inflammation, plasma ADO levels have also been found to be significantly elevated in patients with chronic urticaria compared with healthy controls.^44^ Moreover, animal studies have demonstrated that elevating ADO levels in mice can activate features of chronic diseases, such as pulmonary inflammation and airway remodeling. ^45–47^

Despite its pathophysiological relevance, the ADO-mediated transcriptional program in primary mast cells remains largely unexplored. A search of PubMed Central using keywords related to *adenosine*, *transcriptome*, and *mast cell* identified only one relevant study, which performed transcriptomic profiling of the human mast cell line HMC-1 following ADO receptor stimulation^48^. However, a comprehensive analysis of the ADO-specific transcriptional program in primary peritoneal mast cells (PMCs) is still lacking.

Here, we used bulk RNA-sequencing to characterize the transcriptional programs of PMCs in response to three agonists known to trigger Ca^2+^-dependent mast cell activation, i.e. ADO, Compound 48/80 (C48/80), and antigen stimulation. To characterize the unique ADO-specific response, we identified genes that were differentially expressed exclusively under ADO treatment when compared to both C48/80 and antigen. This analysis identified a distinct set of 393 ADO-regulated genes. Functional analysis using transcription factor activity inference, protein classification, functional enrichment analysis, protein-protein interaction network construction, and topology analysis revealed that these genes are involved in phosphoinositide signaling, vesicle trafficking, glycolysis and mitochondrial activity, and cell cycle arrest. This computational approach not only confirmed consistency with previously described mast cell signaling mechanisms, but also revealed an ADO-specific increase in the expression of genes encoding de novo synthesized mediators, such as *Tgfa* and *Il7*. The regulation of their expression may represent a novel component of ADO-driven mast cell inflammatory responses. Taken together, the identification of an ADO-specific transcriptional program highlights a set of candidate genes for future functional validation in mast cell-mediated processes of tissue inflammation.

## 2 Results

### 2.1 ADO, C48/80, and DNP induce Ca^2+^ release followed by Ca^2+^ influx

To demonstrate that the PMC mast cell model functionally expresses the receptors and downstream signaling molecules necessary for Ca^2+^ dependent activation by ADO, C48/80, and antigens, we used Fura-2 microfluorometry to monitor changes in the intracellular Ca^2+^ concentration ([Ca^2+^]_i_) in PMCs in response to any of three agonists. To separate calcium release from intracellular stores and calcium entry across the plasma membrane, we applied the so-called “re-addition” protocol. In the absence of extracellular Ca^2+^, PMCs responded to all three agonists with a transient elevation of [Ca^2+^]_i_ resulting from calcium release from intracellular stores (Fig. 1A-C). After [Ca^2+^]_i_ returned to the baseline level, extracellular calcium was “re-added” to its physiological concentration of 2 mM. It evoked a fast and prominent [Ca^2+^]_i_ rise due to the agonist-evoked Ca^2+^ entry. All three stimuli: adenosine (10 μM), compound 48/80 (50 μg/mL), and antigen (dinitrophenyl-human serum albumin 100 ng/mL) elicited a reaction with similar pattern of the Ca^2+^ release and Ca^2+^ entry response (Fig. 1A-C). These results demonstrated that all three agonists are able to evoke Ca^2+^-dependent activation pathways in PMCs.

**Figure 1.**
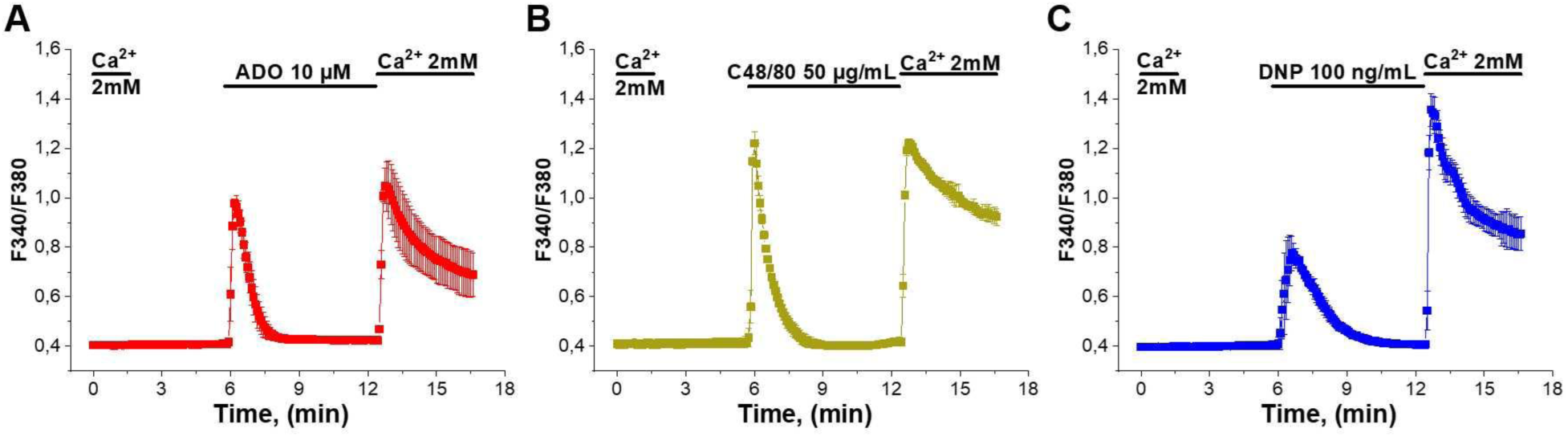
In PMCs, Ca²⁺ release, followed by subsequent Ca²⁺ entry, is induced by ADO, C48/80, and DNP. Fura-2 based calcium imaging of PMCs following treatment with (A) of adenosine (ADO, 10 μM), (B) compound 48/80 (C48/80, 50 μg/mL), and (C) of dinitrophenyl-human serum albumin conjugate (DNP, 100 ng/mL). Changes of [Ca^2+^]_i_ over time are presented as the fluorescence ratio F340/F380 mean values of 3 independent preparations. Error bars represent standard error of mean.

### 2.2 Agonists’ impact on calcium ion channel expression

All three tested agonists evoked calcium signals in PMCs. We then analyzed the expression profile and agonist-evoked transcriptional changes in genes encoding calcium channel constituents and regulators across four stimulation conditions: ADO for 2 hours, Compound 48/80 (C48/80) for 6 hours, and Antigen (DNP) for 2 and 6 hours. A complete summary of the expression levels under all conditions is provided in Table S1. In unstimulated PMCs, we found abundant expression (defined as DESeq2 normalized count > 100) of the following genes encoding channels, subunits and modulators: *Cacnb4*, *Cacng7*, *Ryr3*, *Itpr1*, *Itpr2*, *Itpr3*, *Tpcn1*, *Tpcn2*, *Tmem63a*, *Tmem63b*, *Trpv2*, *Trpm2*, *Trpm4*, *Trpm7*, *Mcoln1*, *Pkd2*, *Orai1*, *Orai2*, *Orai3*, *Stim1*, *Stim2*, *P2rx1*, *P2rx4*, *P2rx7*, *Piezo1*, *Grin2c*, *Grin2d*, and *Calhm2*. Some of these calcium channels, such as Inositol 1,4,5-trisphosphate-gated calcium channel (ITPR1, ITPR2, ITPR3), Transmembrane protein TMEM63A, Transient receptor potential channels TRPV2, Stromal interaction molecule 1 (STIM1), and P2X purinoceptor (P2RX1, P2RX4, P2RX7), were also identified on the protein level in a proteome dataset from unstimulated primary mouse peritoneal mast cells (PMCs).^49^ Furthermore, ITPR1,ITPR2, Two pore segment channel (TPCN1), TMEM63A, TRPV2, STIM1, P2RX1, P2RX4 were identified in proteome dataset of both human skin and fat mast cells.^49^

Among the 11 calcium channel genes differentially regulated by ADO, two were upregulated while nine were downregulated (Table S1). Notably, NMDA receptor *Grin2d* showed the highest ADO-specific log₂ fold change (log_2_FC =-1), representing the most strongly induced transcript. In contrast, transient receptor potential channel *Trpm4* was upregulated in both ADO (2h) and DNP (2h) treatments. A critical molecular regulator of Store-Operated Calcium Entry *Stim1* was consistently downregulated in both ADO (2h) and DNP (2h) conditions. *Tmem63b* encoding for a mechanosensitive cation channel in laminar bodies of AT1 and AT2 cells^50^ displayed upregulation across all agonist treatments, whereas P2X receptor genes *P2rx4* and *P2rx7* were uniformly downregulated across all conditions.

### 2.3 Transcriptional expression of PMC proteases

Given that mast cell proteases are among the most abundantly expressed transcripts, reaching or surpassing the levels of classical housekeeping genes, we examined the expression profiles of major murine mast cell proteases under CONTROL conditions (Fig. S1). *Mcpt1* (mMCP-1) transcripts were undetectable, and *Mcpt2* (mMCP-2) expression remained minimal (Fig. S1). In contrast, *Cma1* (mMCP-5), *Mcpt4* (mMCP-4), *Tpsb2* (mMCP-6), *Tpsab1* (mMCP-7), and *Cpa3* (CPA3) were highly expressed (Fig. S1). These results are consistent with previous findings about proteases expressed in CTMCs^22,23^. Among these, *Cpa3* and *Tpsb2* displayed the highest transcript levels (77,039 ± 11,727 and 82,027 ± 5,376, respectively), followed by *Cma1* (34,382 ± 1,897), *Tpsab1* (11,699 ± 2,514), and *Mcpt4* (7,191 ± 502). These quantitative data demonstrate that *Cpa3* and *Tpsb2* dominate the protease transcriptome in PMCs under basal conditions. In addition, *Ctsg* (cathepsin G) and *Prss34* (mast cell protease 11) were abundantly expressed, aligning with protein-level evidence reported in mouse connective tissue mast cells.^49^

### 2.4 ADO induces a distinct transcriptional response in mast cells

To assess global transcriptional differences among treatment groups, a principal component analysis (PCA) was performed (Fig. S2). The x- and y-axes represent the variance explained by principal components (PC) 1 and 2, respectively. When all samples were analyzed together (Fig. S2A), distinct clustering was observed between the CONTROL and treated groups, with PC1 accounting for 31.44% of the total variance. ADO-, C48/80-, and DNP-treated samples formed separate clusters, indicating treatment-specific transcriptional responses. Treated groups were restricted to ADO (2h) and C48/80 (6h) (Fig. S2B), further demonstrating clear separation from CONTROL samples. Similarly, DNP-treated samples (Fig. S2C) were segregated from CONTROL along PC1, with distinction between 2h and 6h treatments along PC2, suggesting time-dependent transcriptional changes.

To evaluate the transcriptional program induced by ADO, we performed differential expression analysis comparing ADO (2h) to CONTROL and identified 821 upregulated (Fig. 2A) and 630 downregulated genes (Fig. 2B). A notable portion of this response was unique to ADO, as no other stimuli modulated these 223 upregulated and 170 downregulated genes. We visualized the global changes of these ADO-specific genes in a heatmap across all five experimental conditions (seven biological replicates each) (Fig. 2C). The ADO (2h) stimulation induced a visually distinct gene expression pattern compared to CONTROL, which was not seen after C48/80 or DNP stimulation.

We next assessed the overlap with other activation pathways. The ADO (2h) transcriptional profile most closely resembled the 2-hour antigen stimulation (DNP (2h)) response, sharing 548 commonly upregulated (Fig. 2A) and 405 commonly downregulated genes (Fig. 2B). These data demonstrate that ADO drives a unique transcriptional profile, while also sharing a significant number of DEGs with the canonical antigen-mediated activation pathway.

**Figure 2.**
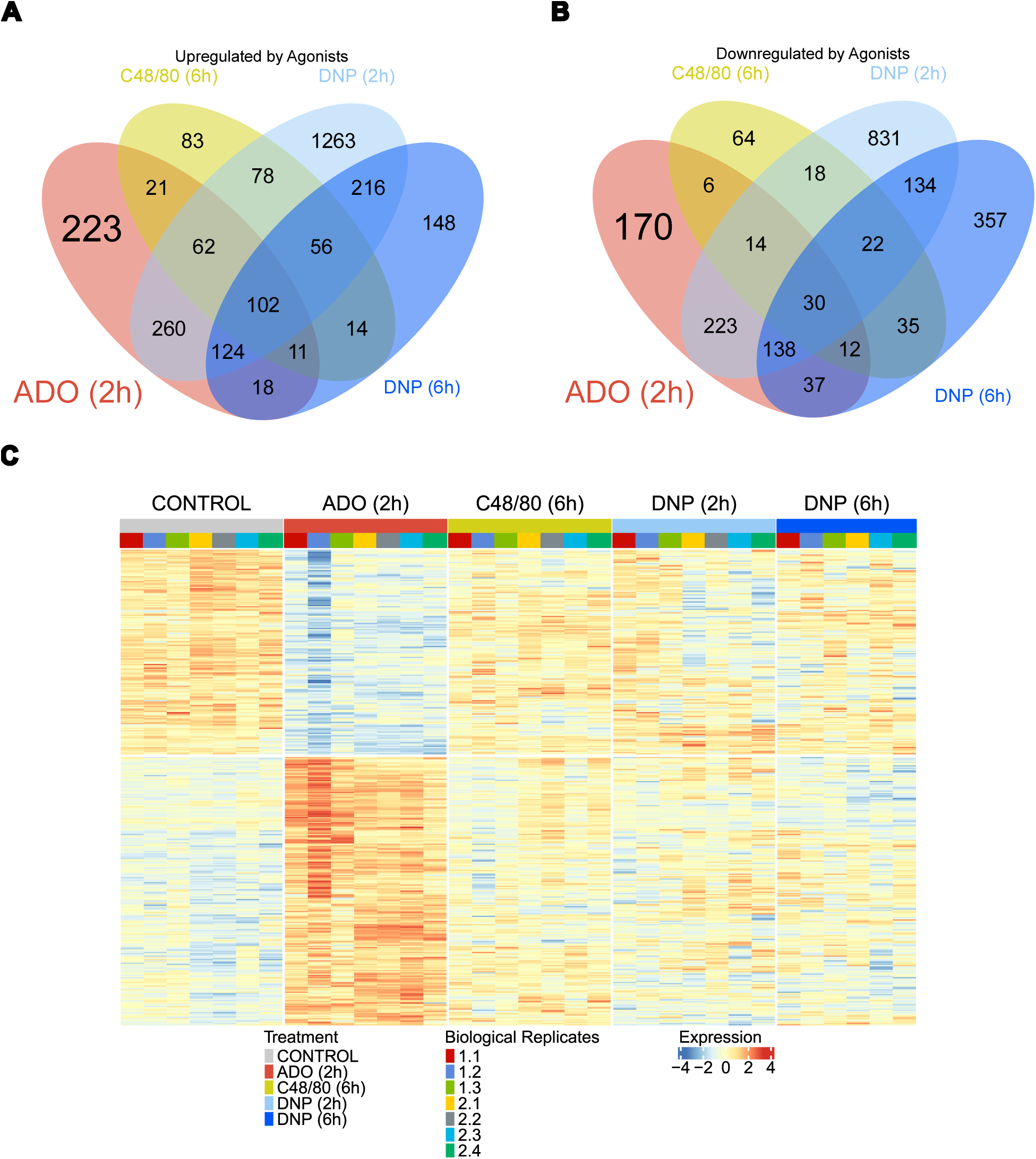
Common and distinct transcriptional responses to mast cell agonists. Venn diagrams showing the overlap of significantly (A) upregulated and (B) downregulated genes across four conditions (ADO (2h, red), C48/80 (6h, yellow), DNP (2h, light blue), and DNP (6h, dark blue)) relative to CONTROL (p_adj_ < 0.01). (C) Heatmap visualizing DESeq2 normalized expression of ADO-specific genes across all five conditions: CONTROL, ADO (2h), C48/80 (6h), DNP (2h) and DNP (6h). Each condition included seven biological replicates, except for the DNP (6 h) treatment, which comprised six replicates due to the unavailability of replicate 2.2.

Overlap with the later DNP (6h) stimulation (255 upregulated, 217 downregulated genes) and the C48/80 (6h) stimulation (196 upregulated, 62 downregulated genes) was also observed (Fig. 2A, B), although to a lesser extent. A core set of 102 genes was upregulated across all four conditions, likely representing a general mast cell activation signature (Fig. 2A). Conversely, only 30 genes were commonly downregulated (Fig. 2B).

To better visualize genes exhibiting both large-magnitude and statistically significant changes in expression, we generated Volcano plots comparing each treatment condition (ADO (2h), C48/80 6 h, DNP (2h), and DNP (6h)) to CONTROL (Fig. S3). These analyses further confirmed distinct transcriptional responses of PMCs to each stimulus.

The differential expression profile of ADO-specific genes is visualized in a Volcano plot (Fig. 3A). Detailed statistics for all ADO-specific protein-coding genes, whose biotype was annotated as “protein-coding” in Ensembl Release 115^51^, are provided in Table S2. The most significantly upregulated ADO-specific genes were involved in cell adhesion, such as the scaffolding protein *Mpp7* and the immunoglobulin superfamily member *Igsf5*, as well as those related to lipid metabolism, including oxysterol-binding protein family *Osbpl6* and prostaglandin transmembrane transporter *Slco3a1*. A strong induction was also observed for signaling molecules, such as the cAMP-responsive element modulator *Crem* and the small GTPases *Rap2b* and *Rab4a*. Other highly induced genes included the apoptosis inhibitor *Niban1*, the pre-mRNA splicing factor *Isy1*, and the calcium-binding protein *Hpcal1*. *Osbpl6, Mpp7*, *Crem*, and *Isy1* were the four most induced genes by ADO with high mean expression across all conditions (DESeq2 normalized counts >100) and the largest absolute log₂ fold changes. They all demonstrated a robust and specific upregulation after two hours of ADO treatment (Fig. 3B). In contrast, a smaller cohort of genes was significantly downregulated, most prominently the growth differentiation factor *Gdf11* and the glutamate receptor subunit *Grin2d* (Fig. 3B).

We next compared our ADO-specific gene signature against a 128 mast cell signature gene set defined by Plum et al. This reference signature was established based on genes exhibiting at least a twofold higher transcript expression in all mast cell populations compared to other analyzed immunocytes^52^. We found an overlap of four upregulated genes: RAS oncogene family member *Rab27b*, latexin *Lxn*, MAS-related GPR family member *Mrgprx2*, Neutral cholesterol ester hydrolase *Nceh1*, and two downregulated genes: NCK-associated protein *Nckap1*, and Endothelin receptor *Ednra*. In particular, the expression of the top 4 most significant ones *Rab27b, Lxn*, *Nckap1*, and *Ednra* was shown in a bar graph (Fig. 3C).

**Figure 3.**
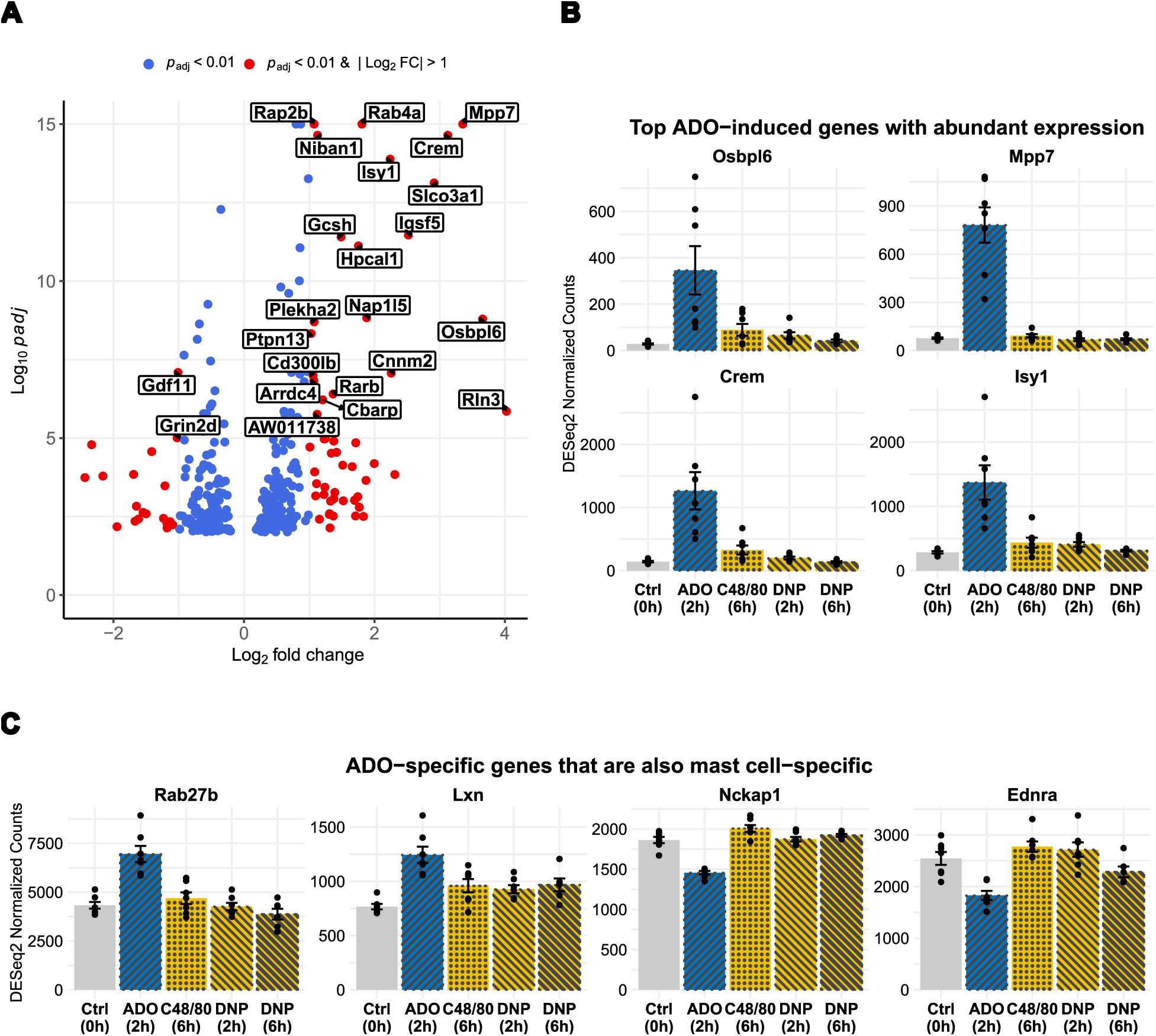
ADO-specific transcriptional program in mast cells. **(A)** Volcano plot of DEGs in mast cells following a 2-hour ADO treatment versus CONTROL. Genes meeting thresholds for both significance (p_adj_ < 0.01) and fold-change (|log_2_FC| > 1) were colored red. Genes meeting only the significance threshold were blue. The most significant DEGs (p_adj_ < 1 × 10−5) were labeled. (B) Normalized expression plots for top four most upregulated and with abundant expression (DESeq2 normalized counts > 100) ADO-specific DEGs across five conditions: CONTROL, ADO (2h), C48/80 (6h), DNP (2h), and DNP (6h). Each condition included seven biological replicates, except for the DNP (6 h) treatment, which comprised six replicates due to the unavailability of replicate 2.2. **(C)** Normalized expression plots for ADO-specific DEGs that are also tissue-resident mast cell-specific; all other conditions are as described in (B).

### 2.5 Integrated functional and regulatory enrichment analysis of ADO-specific genes

To estimate the ADO-evoked activation of transcription factors (TFs), transcription factor activity inference was performed. This analysis revealed significant modulation of two TFs: *E2f1* and *Ctnnb1*, whose target gene expression was visualized in Fig. 4A and 4B, respectively.

**Figure 4.**
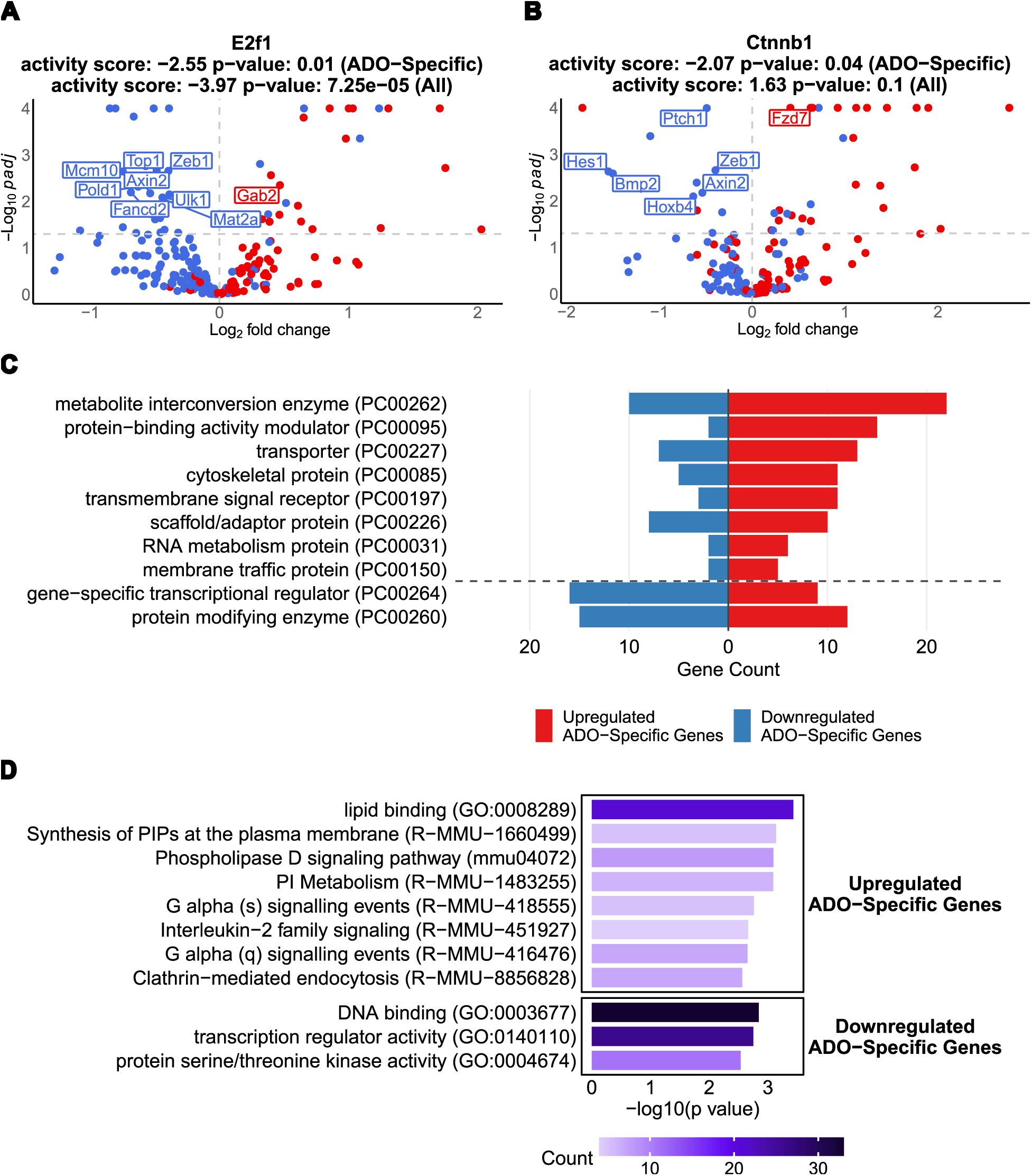
Integrated functional and regulatory enrichment analysis of ADO-specific genes. **(A, B)** Volcano plots displaying the inferred activity of TFs based on their ADO-specific target genes, as determined by ULM analysis. Labeled targets are ADO-specific genes. **(C)** A bar graph showing the top ten annotated protein classes in upregulated and downregulated ADO-specific genes according to PANTHER classification system. **(D)** Bar graph showing significantly enriched pathways from GO Molecular Function, KEGG, and Reactome databases that satisfy *p*-value < 3 × 10^−03^. The gene count within each term is visualized by color density.

The TF *Ctnnb1* encodes β-catenin, a central component of the Wnt signaling pathway. Although *Ctnnb1* activity was not significantly altered across all target genes (activity score = 1.63, p = 0.1), its activity was significantly suppressed across the ADO-specific gene set (activity score = −2.07, p = 0.04) by ADO treatment. This indicates that ADO engages a dedicated pathway to suppress the transcriptional output of the canonical Wnt signaling pathway. *E2f1* is a transcription factor that controls the cell cycle. Its activity was strongly and globally inhibited across both the full target set (activity score = −3.97, p = 7.25 × 10^−5^) and the ADO-specific subset (activity score = −2.55, p = 0.01), as evidenced by coordinated suppression of DNA synthesis genes such as *Mcm10*, *Pold1*, *Top1*, and *Fancd2*.

Gene annotation via the Panther classification system identified protein classes for 137 of 184 upregulated and 85 of 136 downregulated ADO-specific genes. The classes with at least 5 annotated proteins were listed in a bar graph (Fig. 4C). The most enriched protein class was metabolite interconversion enzyme (PC00262), which included a broad set of genes essential for core metabolic processes. Key examples include enzymes involved in glycolysis (*Aldoa*, *Pfkp*), fatty acid metabolism (*Elovl5*, *Cpt1a*), and mitochondrial energy production (*Impdh1*, *Sdhb*, *Ldha*) (Table S3).

The second major class, protein-binding activity modulators (PC00095), suggests widespread activation of intracellular signaling networks. This is highlighted by the upregulation of numerous small GTPases (*Arf2*, *Rab27b*, *Rhoq*, *Rap2b*, *Rab11fip5*, and *Rab4a*) and their regulators, such as the guanyl-nucleotide exchange factors *Trio* and *Cyth1*, and GTPase-activating protein *Arhgap25* and *Sipa1l1* (Table S3). The prominence of GTPase-related signaling also drew our attention to the unique upregulation of GTPases *Gvin1* and *Gvin2*, which encode GTPase, very large interferon inducible 1 and GTPase, very large interferon inducible 2, following ADO treatment (Table S3). The third class, transporters (PC00227), shows an increase in genes involved in the movement of molecules across cellular membranes. This includes components of the ATP synthase complex (*Atp5f1d*, *Atp5mk*), various ion channels (*Cbarp*, *Kcnn4*, *Vdac3*), and multiple solute carrier organic anion transporter family members like *Slco3a1*, *Slc20a2*, and *Slc16a3* (Table S3). In the enrichment of cytoskeletal protein (PC00085), we observed an upregulation of genes involved in both the actin and microtubule networks (Table S3). Enrichment was also observed for the transmembrane signal receptor class (PC00197), which included G-protein coupled receptors (*Grm5*, *Lpar2*, *Mrgprx2*), the pattern recognition receptor *Tlr4*, Frizzled family members *Fzd6* and *Fzd7*, and cytokine receptor subunits *Csf2rb* and *Csf2rb2* (Table S3). The scaffold/adaptor protein class (PC00226) was also represented, including the highly upregulated *Mpp7*, as well as signaling regulators such as *Ywhaz* and *Arrb1* (Table S3).

In contrast to the upregulation of metabolic and structural genes, a strong trend of downregulation was observed in two key regulatory classes: Gene-Specific Transcriptional Regulators (PC00264) and Protein Modifying Enzymes (PC00260) (Fig. 4C). The first class contains many transcription factors of gene expression programs. This includes a large family of C2H2 zinc-finger proteins (*Glis2*, *Zeb1*, *Ikzf2*) as well as key developmental factors such as *Gata1*, *Gata2*, and *Hoxb4* (Table S4). This widespread downregulation suggests a major shift or shutdown of established transcriptional programs. The second class contains genes regulating protein function through post-translational modifications. This is highlighted by the decreased expression of numerous non-receptor serine/threonine protein kinases (*Ksr1*, *Taok3*, *Prkcd*) and components of the ubiquitin system, including multiple E3 ubiquitin-protein ligases (*Pias2*, *Itch*, *Trim58*) (Table S4). This points to a significant alteration in cellular signaling pathways and protein degradation.

To further understand the molecular functions and signaling pathways involved, we performed functional enrichment analysis of genes specifically upregulated or downregulated by ADO treatment using databases from GO, KEGG, and Reactome. The analysis revealed a diverse set of functions, with ADO promoting pathways involved in cell signaling while suppressing those involved in nuclear regulation and gene expression (Fig. 4D).

Among the upregulated genes, the most significant enrichment was found for pathways associated with membrane signaling and lipid metabolism. The top enriched terms included lipid binding (GO:0008289), Synthesis of PIPs at the plasma membrane (R-MMU-1660499), the Phospholipase D signaling pathway (mmu04072), and PI Metabolism (R-MMU-1483255) (Fig. 4D). Furthermore, multiple G-protein signaling pathways, such as G alpha (s) and G alpha (q) signaling events, were significantly enriched, indicating a broad activation of signal transduction cascades originating at the cell membrane. In contrast, the most prominent downregulated categories included DNA binding (GO:0003677), transcription regulator activity (GO:0140110), and protein-modifying enzymes (PC00260) (Fig. 4D), reflecting a broad suppression of genes involved in transcriptional regulation, signaling, and post-translational modification in response to ADO stimulation.

### 2.6 Protein-protein interaction network revealing ADO-evoked changes in key hubs of metabolism and signaling

To further investigate the functional relationships between the ADO-specific genes, we constructed a protein-protein interaction network. This analysis identified distinct, highly interconnected modules corresponding to core metabolic processes and phosphatidylinositol signaling, highlighting key hub genes that likely orchestrate these cellular responses (Fig. 5).

Four of the top seven hub genes, identified by the MNC algorithm, were central components of the metabolic process network. Notably, Lactate dehydrogenase A (*Ldha*) and Aldolase A (*Aldoa*) are key enzymes in glycolysis. In contrast, the iron-sulfur subunit B of the succinate dehydrogenase complex (*Sdhb*) and the F1 subunit delta of ATP synthase (*Atp5f1d*) are integral to mitochondrial function. A second central module was organized around Phosphatidylinositol-Related signaling. This network connected numerous phospholipases, kinases, and their associated proteins, indicating a comprehensive remodeling of this critical second messenger pathway. Three hub genes were central to this module: Phospholipase C gamma 1 (*Plcg1*), Phospholipase C beta 3 (*Plcb3*), and Phosphatidylinositol-4,5-bisphosphate 3-kinase catalytic subunit beta (*Pik3cb*) (Fig. 5). Unlike the metabolic module, this signaling network comprised a mix of both upregulated and downregulated genes, suggesting a complex, fine-tuned regulation of phosphoinositide signaling. In summary, the network analysis pinpoints critical hub genes that connect ADO stimulation to two primary cellular outcomes: a coordinated upregulation of central energy metabolism and a complex remodeling of the phosphatidylinositol signaling cascade.

**Figure 5.**
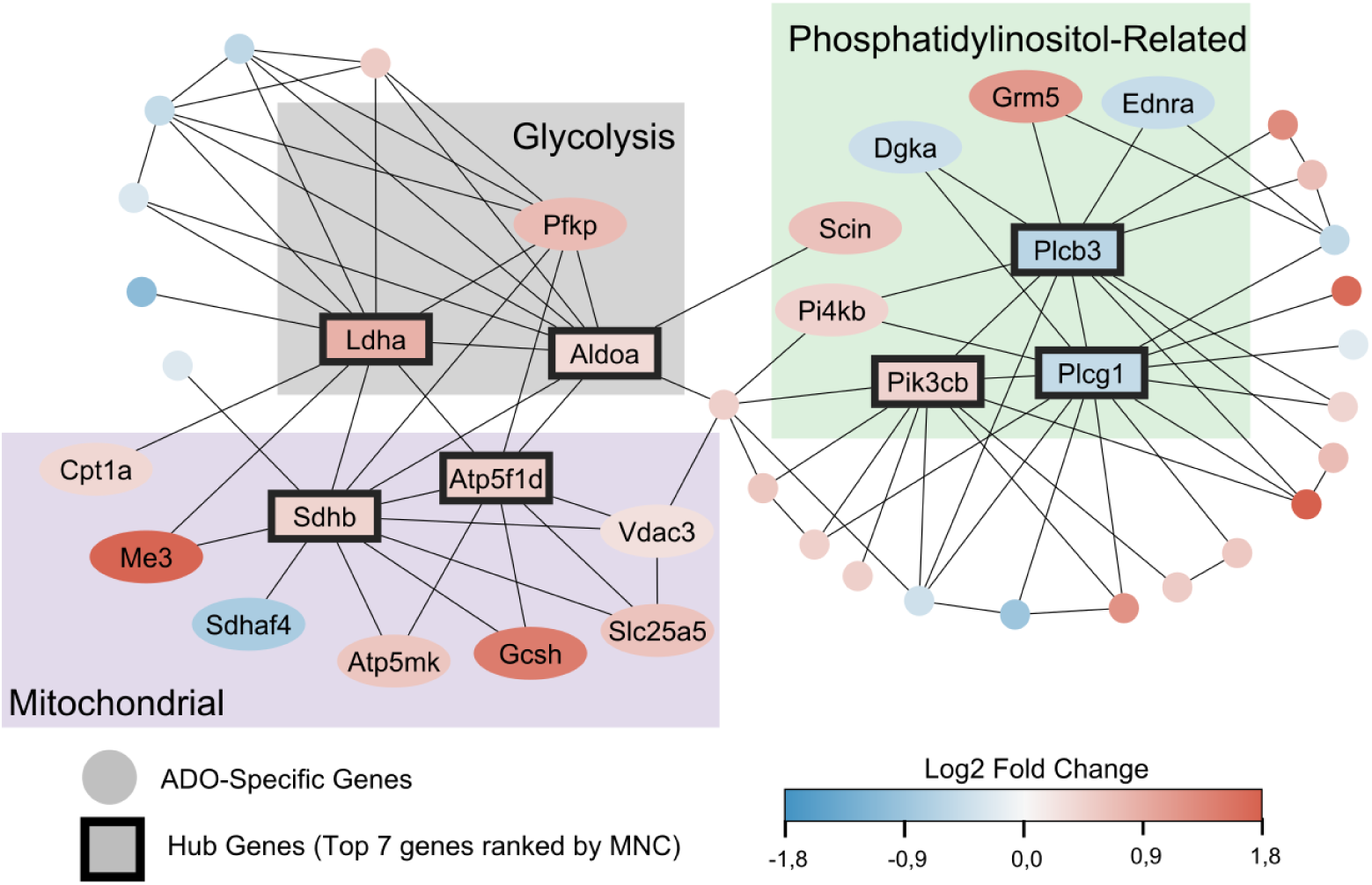
Protein-protein interaction network analysis of ADO-regulated genes and identification of hub genes. The network illustrates the predicted physical and functional interactions among genes whose expression is specifically regulated by ADO. The color of each node (gene) corresponds to its log_2_FC: with red indicating upregulation and blue indicating downregulation. Seven highly connected hub genes, outlined in black, are central to the network’s function. These hub genes are grouped into three key functional modules indicated by their respective node background colors: Glycolysis (Grey), Mitochondrial Activity (Purple), and Phosphatidylinositol-Related Activity (Green).

### 2.7 Topology analysis revealing a causal structure of key ADO-modulated pathways

To elucidate the causal regulatory architecture of pathways significantly affected by ADO, we performed topology analysis on Reactome pathways using SEMgraph. We identified six significantly modulated regulatory networks, each involving at least three ADO-specific genes, highlighting key activation events in metabolism, endocytosis, and nuclear processes (Fig. 6).

Among these, Clathrin-mediated endocytosis exhibited the best model fit (*dev*/*df* = 2.05, srmr = 0.08) (Fig. 6A). This network involved a mixed expression profile, with *Tgfa* and *Arrb1* being significantly upregulated, while *Fcho1* was repressed. The phosphatidylinositol metabolism pathway was also significantly perturbed (pNodeAct = 3.43*e*^−14^ , pNodeInh = 9.80*e*^−10^), implicating key upstream kinases *Pik3cb*, *Pi4kb*, and *Pip5k1c* that regulate downstream effectors such as *Plcg1*, *Adgpk*, and *Rab4a* (Fig. 6B). Two central metabolic pathways, the TCA cycle and respiratory electron transport (pNodeAct = 9.21*e*^−10^) and Glycolysis (pNodeAct = 3.24*e*^−09^), were significantly activated, driven by the upregulation of enzymes like *Sdhb*, *Ldha*, *Pfkp*, and *Aldoa* (Fig. 6C, 6D). Conversely, the SUMOylation pathway exhibited strong inhibition (pNodeInh = 1.58*e*^−08^) (Fig. 6E). Finally, the Nuclear Envelope Breakdown pathway showed bidirectional regulation, with *Emd* and *Lmnb1* being activated and *Nup153* being repressed (Fig. 6F). In summary, this topological analysis provides a mechanistic framework for understanding how ADO orchestrates mast cell responses through the activation of metabolic and signaling pathways and the modulation of nuclear and post-translational regulatory circuits.

### 2.8 ADO’s impact on the expression of genes involved in the release of de novo synthesized mediators

To identify ADO’s effect on de novo synthesized mediators in mast cells, we curated a list of genes encoding lipid mediators, cytokines, chemokines, growth factors, and their receptors (see Table S5). We then checked the expression patterns of genes that were either uniquely responsive to ADO or significantly modulated by ADO treatment (p_adj_ < 0.01, Fig. S4).

Notably, *Pla2g4a* and *Dagla*, key enzymes responsible for liberating arachidonic acid and 2-Arachidonoylglycerol from membrane phospholipids, were potently upregulated following ADO stimulation. Additionally, the growth factor *Tgfa* and the cytokine *Il7* were upregulated in response to ADO. Among ADO-specific immune receptors, we found upregulation of the pattern recognition receptor *Tlr4*, the Frizzled family members *Fzd6* and *Fzd7*, and the cytokine receptor subunits *Csf2rb* and *Csf2rb2*, and downregulation of the endothelin-1 receptor *Ednra*.

**Figure 6.**
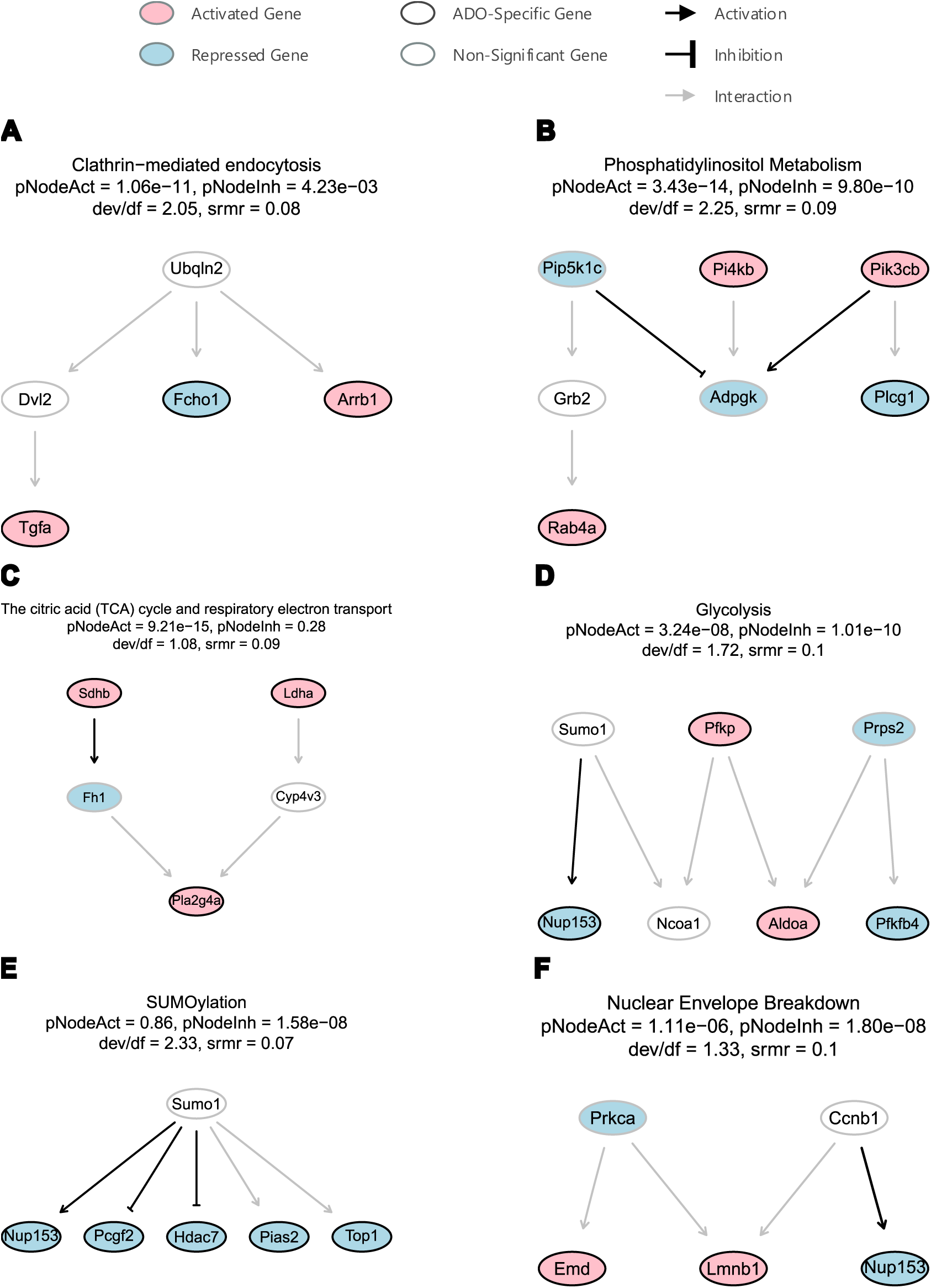
Topological analysis of Reactome pathways seeded with ADO-specific genes. Six significant regulatory networks were identified from the Reactome database using SEMgraph. Networks shown were selected for containing at least three ADO-specific genes and meeting model fit criteria (SRMR < 0.1 ; deviance/df < 3). The combined effect of all nodes was considered significantly activated if *pNodeAct* < 0.05 and/or inhibited if *pNodeInh* < 0.05.

## 3 Discussion

ADO is a mediator implicated in a variety of inflammatory processes, and it also plays an essential role in mast cell activation and its fine-tuning. Since ADO alone can’t induce massive mast cell degranulation and typically enhances the release of mediators evoked by immunological stimuli, studying ADO-stimulated transcriptional responses could provide new insights into the intracellular signaling pathways triggered by this agonist. The number of reports on this is very limited and includes only studies using immortalized mast cell lines.

In most of studied mast cell models, ADO stimulation evokes elevation of intracellular Ca^2+^ concentration. Intracellular Ca^2+^ is a ubiquitous and essential trigger of specific functions of eukaryotic.^53^ For example, elevation of [Ca^2+^]_i_ in mast cells activates not only the exocytosis but also the transcription factor NFAT, which regulates gene expression.^54,55^ Mast cell degranulation is indispensable to [Ca^2+^]_i_ elevation. In PMCs, the stimulation of FcεRI, Mrgprb2, and ADO receptors evoked Ca^2+^-release followed by Ca^2+^-entry is of similar pattern for all three stimuli. Thus, in our mast cell model used in this study, ADO triggered functional responses that were principally similar to those evoked by other mast cell activators.

### ADO’s effects on calcium channel expression

Transcriptomic analysis provides a complementary approach for understanding how PMCs regulate channel abundance and their regulators, thereby determining calcium homeostasis during agonist stimulation over time at the transcriptional level. Among Ca²⁺ channel-related genes, only *Trpm4* and *Tmem63b* were upregulated upon ADO stimulation. Our previous work identified TRPM4 as a critical limiting factor for antigen-evoked calcium rise in mouse PMCs, in which loss of TRPM4 leads to exaggerated calcium influx and enhanced degranulation.^56^ *Tmem63b* (also known as *OCaR2*), a mechanosensitive cation channel, was identified to be upregulated by 7-fold after LPS treatment in bone marrow-derived mast cells^57^, suggesting that it may participate in mast cell activation. In our study, *Tmem63b* expression increased modestly (1.4-fold, 1.6-fold, 1.5-fold, and 1.9-fold) in response to ADO (2 h), C48/80 (6 h), DNP (2 h), and DNP (6 h), respectively. Conversely, the downregulation of *Grin2d*, *Itpr1*, *Stim1*, *P2rx4*, and *P2rx7* may indicate the engagement of a transcriptional negative-feedback mechanism to prevent excessive calcium entry and maintain calcium homeostasis in PMCs following ADO stimulation.

### ADO-specific transcriptional responses

Our analysis reveals that ADO induces a specific transcriptional program in mast cells. Namely, while a majority (65%) of ADO-induced genes overlap with the canonical antigen-triggered response, a significant portion (27%) is unique to ADO stimulation. The major shared signature with antigen is comprehensible since ADO is known to augment the canonical FcεRI-induced degranulation.^32^ In particular, we observed significant upregulation of *Hdc* and *Tpsab1* following both ADO (2h) and DNP (2h) stimulation. *Hdc* encodes histidine decarboxylase, the rate-limiting enzyme responsible for histamine synthesis, while *Tpsab1* encodes the tryptase mMCP-7, a major protease stored in mast cell granules. Notably, the upregulation of *Hdc* that we observed in PMCs is consistent with findings from a previous transcriptomic study of the ADO-treated human mast cell line HMC-1^58^. On the other hand, the unique ADO-induced molecular signature is of particular interest to us. It could provide an insight into the unique ADO signaling pathway that contributes to the release of inflammatory mediators without massive degranulation. For example, we found *Hdac7* downregulation specifically in ADO. According to the CollecTRI database^59^, *Hdac7* negatively regulates *Hdc* expression, suggesting an ADO-specific pathway regulating histamine synthesis.

### ADO’s effects on the expression of molecules determining GPCR signaling

ADO exerts its effects primarily by activating ADO receptors on the plasma membrane. Upon stimulation, these receptors signal their respective G proteins, initiating downstream signaling cascades.^29^ In line with this, pathway enrichment analysis of ADO-responsive genes demonstrated significant enrichment for both G alpha (s) and G alpha (q) signaling events.

Activation of G alpha (s) protein increases AC activity and cAMP production.^29^ The significant upregulation (∼9-fold) of *Crem* (cAMP-responsive element modulator) gene in our data suggests the activation of a canonical GPCR-cAMP signaling axis.^60^ This is consistent with the previous findings that the human gene CREM is significantly upregulated upon ADO receptor activation in human mast cell line HMC-1,^48^ and our study further demonstrates that the transcriptional response of *Crem* is unique to ADO. Concurrently, activation of G alpha (q) protein leads to PLC activation.^61^ Our PPI analysis identified *Plcb3* (Phospholipase C β3) and *Plcg1* (Phospholipase C, gamma 1) as hub genes in the ADO-specific response network, and both were downregulated. This highlights their regulatory role and suggests a transcriptional negative feedback loop after activation.

### ADO’s effects on the PI3K/Akt signaling axis

Another hub gene, *Pik3cb* (Phosphatidylinositol-4,5-bisphosphate 3-kinase catalytic subunit beta), was upregulated in response to ADO. A previous study in RBL-2H3 cells indicated that ADO activates Gi-coupled A3 receptors, leading to protection against apoptosis via a pathway involving the Gi_βγ_ subunits, PI3Kβ, and protein kinase B (Akt).^62^ Consistent with this, our data show that G gamma subunits *Gng4* and *Pik3cb,* and their docking protein *Gab2,* were uniquely upregulated in ADO, supporting an enhancement of the PI3K/Akt signaling axis and suggesting a pathway involved in promoting mast cell survival and metabolic activity.

### ADO’s effects on the expression of molecules determining vesicle trafficking

Our protein classification analysis revealed ADO-specific upregulation of genes associated with vesicle trafficking, including small GTPases (e.g., *Rab27b*, *Rab4a*), cytoskeletal-associated proteins (e.g., *Myo5a*, *Arpc3*), and scaffold/adaptor molecules (e.g., *Arrb1*, *Ywhaz*). Among these, *Rab27b* plays a crucial role in mast cell degranulation, particularly by regulating the transition of vesicle transport from microtubule-to actin-based motility.^63^ *Myo5a*, encoding the motor protein MYO5A, is part of the RAB27a–Mlph–MYO5A complex, which regulates distinct steps in the BMMC degranulation pathway.^64^ Likewise, *Rab4a* has been identified as a key regulator of mast cell exocytosis through vesicle trafficking pathways.^65^ In addition to exocytic machinery, enrichment of clathrin-mediated endocytosis pathways was observed, suggesting active internalization and recycling of receptors or membrane components during ongoing vesicle turnover and signaling.^66^ *Arrb1*, encoding β-arrestin 1, functions as a molecular scaffold linking G-protein-coupled receptors to clathrin-mediated endocytosis.^67^ Notably, β-arrestin 1 mediates agonist-dependent internalization and desensitization of the MRGPRX2 receptor in human mast cells.^68^ Moreover, growth factor *Tgfa*, which was uniquely upregulated by ADO treatment, undergoes endocytic trafficking via a clathrin-dependent pathway.^69^ Collectively, these findings highlight that ADO stimulation reprograms the vesicular trafficking machinery in PMCs, which may contribute to the potentiating effect of ADO on antigen-evoked degranulation.

### ADO’s effects on the expression of inflammatory mediators

We observed an increased expression of a range of lipid-mediator-producing enzymes, cytokines, chemokines, and immune receptors in response to ADO. Notably, enzymes involved in the liberation of precursor fatty acids from membrane phospholipids were upregulated, such as *Pla2g4a* and *Dagla*. Additionally, the growth factor *Tgfa* and the cytokine *Il7* were significantly upregulated in response to ADO. In contrast, microarray analysis using HMC-1 cells reported a twofold downregulation of IL7 following Cl-IB-MECA treatment.^70^ We also observed a unique upregulation of immune receptors, including *Csf2rb*, *Csf2rb2*, and *Tlr4*, possibly reflecting a feed-forward mechanism that sensitizes the cell to the newly produced cytokines.

### ADO-dependent transcriptional effects in metabolism and cell cycle

Glycolysis and mitochondria are critical for mast cell activation since the inhibition of glycolysis and ATP production attenuated IL-33-mediated and Lipopolysaccharide-induced mast cell function.^71,72^ Through PPI and topology analysis, we also identified activated core metabolic pathways, including glycolysis and mitochondrial activity. In addition, our analysis revealed an ADO-specific suppression of signaling pathways associated with cell proliferation. Our TF activity analysis indicated a reduced activity of *Ctnnb1* (β-catenin) and *E2f1*. Dysregulated *β*-catenin signaling has been shown to promote the expansion of bone marrow-derived connective tissue-type mast cells, systemic inflammation, and colon cancer.^73^ Furthermore, the Wnt/β-catenin pathway is a well-established driver of cell proliferation.^74,75^ E2F1, in turn, is a key regulator of G1/S transition and DNA synthesis, driving expression of genes required for S-phase entry.^76^ Notably, E2F1 is a novel TF regulating Ctnnb1 expression.^77^ The suppression of both TFs suggests a shutdown of the proliferation-promoting Wnt/β-catenin-E2F axis and indicates cell cycle arrest under ADO stimulation.

We also noted a significant topological pathway of nuclear envelope dynamics involving upregulation of *Lmnb1* (Lamin B1) and *Emd* (Emerin) and downregulation of *Nup153*. Upregulation of *Lmnb1* and *Emd* suggests reinforcement of the nuclear lamina, potentially increasing its rigidity and resistance to remodeling^78–80^, while downregulation of *Nup153*, a key component of the nuclear pore complex required for pore disassembly during mitosis, may hinder nuclear envelope breakdown and thereby limit cell cycle progression.^81^ Together, these changes suggest reduced nuclear plasticity and cell-cycle arrest specific to ADO treatment.

### Limitations of our study

While transcriptomic analysis provides a comprehensive framework for hypothesis generation, it has limitations. First, mRNA abundance does not always correlate with downstream protein expression or enzyme activity. It will be essential to validate these findings at the protein level (e.g., proteomic approach) and by measuring functional outputs (e.g., inflammatory mediators release, cAMP, lactate production, oxygen consumption rate, ATP levels, degranulation, cell proliferation, etc.), which should be a matter of our future studies. Second, because calcium signals exhibit spatial and temporal dynamic^53^, our study could not distinguish transcriptional responses driven by calcium release from those triggered by calcium influx. This limitation could be addressed in future work by using pharmacological blockers or genetic models to delineate the specific contribution of defined calcium signals to the observed transcriptional changes. Finally, calcium-independent signaling pathways that influence gene expression also warrant further investigation.

## 4 Conclusions

Our study reveals that ADO (compared with activators of FcεRI and Mrgprb2 receptors) alone can elicit a distinct mast cell activation program characterized by transcriptional remodeling of cellular/intracellular signaling, metabolic, and vesicular pathways. This response may promote the synthesis and release of specific inflammatory mediators, as well as the synthesis of components of the exocytosis machinery. Future studies should validate these ADO-driven mechanisms at the protein level, assess their associated functional responses, and further investigate the transcriptional programs triggered by ADO-specific defined calcium signaling.

## 5 Materials and methods

### 5.1 Mice

All procedures on animals were performed in accordance with the approval of the federal state of Baden-Württemberg, Germany. Mice were bred and maintained at the central animal facility of the University of Heidelberg under specific pathogen-free conditions. They were provided with drinking water and food *ad libitum*. Mice were killed by a lethal dose of CO_2_. The mice used for isolation of PMC mast cells in this study were control mice with the genotype Orai1^flox/flox^; Orai2^flox/flox^ ^82,83^ of the C57Bl6/N genetic background (backcrossed with C57Bl/6N mouse strain obtained from *Charles River, USA* at least 8 generations). Adult 9-12 weeks old male mice were used for the experimental procedures.

### 5.2 Peritoneal mast cells primary culture and agonist stimulation

PMCs were isolated and cultured as previously described^84^. Briefly, the cells were obtained by peritoneal lavage with RPMI medium, centrifuged, and subsequently resuspended in culture medium (4 mL per mouse). The RPMI-based culture medium additionally contained 20% fetal calf serum (FCS), 1% Penicillin-Streptomycin, 10 ng/mL of IL-3 and 30 ng/mL of Stem Cell Factor. For one preparation, the cells isolated from 2-3 mice were pooled. The isolated cell suspension was cultured at 37^∘^C and 5% CO_2_. Two days after the isolation, the culture medium was changed, and all non-adherent cells were removed by aspiration. On day nine, the cells were split and cultivated further at a concentration 1 × 10^6^of cells/mL. The PMCs were used for the experiments 12-16 days after isolation. For the antigen stimulation experiments, the cells were treated overnight with 300 ng/mL anti-Dinitrophenyl IgE antibodies. The PMCs for RNA sequencing were stimulated in the RPMI-based culture medium at a concentration 2 x 10^5^ cells/ml at 37^∘^C and 5% CO_2_.

### 5.3 Microfluorimetric intracellular free Ca^2+^ concentration measurements

PMCs were loaded with Fura-2 by incubating them in a physiological salt solution containing 2.5 μM Fura-2 AM and 0.1% Pluronic F-127 for 30 min at room temperature on the coverslips covered with concanavalin A (0.1 mg/mL) for cell immobilisation. The intracellular free Ca^2+^ concentration was measured on the stage of AxioObserver-A3 inverted microscope (Zeiss, Germany) equipped with a 40x (1.3NA) immersion oil objective (Zeiss, Germany) in a perfused 0.5 mL chamber in Physiological Salt Solution at room temperature. The Physiological Salt Solution (PSS) contained (in mM): NaCl 135, KCl 6, CaCl_2_ 2, MgCl_2_ 1.2, glucose 12 and HEPES 10; pH 7.4 (NaOH). Nominally Ca^2+^-free PSS was made by excluding CaCl_2_ salt from the composition. At the beginning of each experiment, the cells were washed thoroughly with PSS. The fluorescence signal was obtained by alternately exciting the Fura-2 with light of 340 nm and 380 nm wavelength (50 ms exposure time) using a pE-800fura (CoolLED, United Kingdom). Emitted fluorescent signal was filtered at > 510 nm and detected by a charged-coupled device camera AxioCam MRm (Zeiss, Germany). The fluorescent ratio F_340_/F_380_ was measured with an acquisition rate of 5 s per cycle. Both the camera and the Light source were controlled by the Zen 3.2 software (Zeiss, Germany), allowing for the recording of fluorescent signal intensities in particular cell-attributed Regions of Interest (ROIs).

### 5.4 Information about samples undergoing RNA sequencing

A total of seven independent PMC cell preparations were generated, each from 2 or 3 mice. Each PMC preparation was stimulated with four experimental conditions: ADO (10 µM) for 2 hours (ADO (2h)), C48/80 (50µg/mL) for 6 hours (C48/80 (6h)), DNP (100 ng/mL) for 2 hours (DNP (2h)), and DNP (100 ng/mL) for 6 hours (DNP (6h)). From each PMC preparation two technical replicates with vehicle control treatment (CONTROL) were further processed. RNA isolation and subsequent sequencing were performed in two batches. The corresponding samples were termed 1.1, 1.2 and 1.3 (Batch 1), and 2.1, 2.2, 2.3 and 2.4 (Batch 2). For the second biological replicate from batch 2 (2.2), the data for the DNP 6-hour-treated condition were unavailable for this analysis, so six independent biological replicates were analyzed for this condition.

For RNA isolation, each sample (0.2 – 2 x 10^5^ PMCs) was lysed in 400 µL RLT Buffer supplemented with 4 μL-Mercaptoethanol. Isolation of mRNA was performed using RNA Micro KIT (Qiagen 56304); at the end mRNA was eluted in 14 μL buffer and 10 µL RNA of each sample was utilized for the further transcript library synthesis. Transcript libraries were prepared using “Ovation SoLo RNA-Seq Library Preparation Kit” (NuGEN) and 25 ng of the labeled cDNA per sample was utilized for the further sequencing. For the deep sequencing a “NextSeq 2000“ (Illumina) sequencing platform was used (EMBL Heidelberg Genomics Core Facility). The sequencing reads included three types: 75 bp (single-end) for sample 1.1, 61 bp (paired-end) for samples 1.2 and 1.3, and 100 bp (paired-end) for samples 2.1-2.4. For each sample, at least 20 million reads were obtained. The ENSEMBL mouse genome database was used for the transcript identification.

### 5.5 Bioinformatic analysis of RNA-Seq data

#### 5.5.1 Preprocessing and quantification

Raw RNA-Seq reads were processed using the nf-core/rnaseq pipeline (version 3.14.0)^85^. The pipeline was executed using Nextflow^86^ with the Singularity profile^87^ to ensure reproducibility via containerized environments. Read alignment was performed against the *Mus musculus* GRCm39 reference genome (GRCm39.primary_assembly.genome.fa) using STAR (version 2.7.11a)^88^. Gene-level quantification was carried out using RSEM (version 1.3.1)^89^ with annotations from GENCODE release M35 (gencode.vM35.primary_assembly.annotation.gtf).

#### 5.5.2 Differential expression analysis

Differential gene expression analysis was conducted using the DESeq2 package (version 1.42.0)^90^ in R. Prior to analysis, gene counts were filtered to exclude genes with low expression; specifically, only genes exhibiting a raw count of at least 10 in a minimum of 7 samples were retained. This threshold ensures that analyzed genes demonstrate substantive expression in at least one experimental condition, considering the seven biological replicates per condition.

Technical replicates for CONTROL samples were merged by summing their counts using the collapseReplicates function in DESeq2 to increase sequencing depth for those biological samples. The experimental design formula, ∼ batch + treatment, was employed to model and account for potential batch effects arising from batch 1 and batch 2 data, thereby enhancing the sensitivity for detecting treatment-specific expression changes.

The filtered count matrix input to DESeq2 contained 16,177 genes and 34 samples (after merging technical replicates and accounting for the missing DNP (6h) sample). Differentially Expressed Genes (DEGs) between conditions were identified using the Wald test. The resulting *p*-values were adjusted for multiple comparisons using the Benjamini-Hochberg (BH) procedure. A gene was considered significantly differentially expressed if its BH-adjusted *p*-value (p_adj_) was less than 0.01.

#### 5.5.3 Identification and visualization of ADO-specific genes

To identify genes specifically modulated by ADO treatment, DEGs from four comparisons (ADO (2h) vs CONTROL, C48/80 (6h) vs CONTROL, DNP (2h) vs CONTROL, and DNP (6h) vs CONTROL) were compared using Venn diagrams. Genes significantly up-or downregulated exclusively in the ADO (2h) vs CONTROL comparison, but not in the other three comparisons, were classified as ADO-specific DEGs. A heatmap was generated to display the expression profiles of these ADO-specific genes across all five conditions.

Subsequently, a volcano plot was utilized to visualize ADO-specific genes with absolute value of log_2_ Fold Change (|log_2_FC|) > 1 and p_adj_ < 0.01. Moreover, eight representative candidate genes were selected and shown via bar graphs. For visualization, expression data were batch-adjusted using ComBat-seq^91^ and normalized using size factors from DESeq2.

#### 5.5.4 Transcription factor activity inference

Transcription factor (TF) activity inference was conducted using the decoupleR package (version 2.9.1)^92^. The input comprised stat, p_adj_ , and log _2_ fold change values derived from DESeq2 differential expression analysis. TF-target associations were sourced from the CollecTRI database^59^, a comprehensive collection of TF-target relationships aggregated from 12 distinct resources. TF activities were estimated using the default method in decoupleR: the univariate linear model (ULM). TFs were considered significantly activated or repressed if they exhibited an absolute activity score (|score|) > 2 and a *p*-value < 0.05. To identify TFs specifically associated with the ADO response, TF activity inference was performed on ADO-specific genes (see section 2.5.4), and the resulting significantly regulated TFs were considered ADO-specific TFs.

#### 5.5.5 Protein classes classification and functional enrichment analysis

Protein classes classification was conducted using PANTHER (protein analysis through evolutionary relationships) classification system (version 19.0)^93^. The input consisted of 393 ADO-specific DEGs (annotated with Ensembl IDs).

Pathway over-representation analysis was conducted using clusterProfiler^94^ and ReactomePA^95^. Enrichments were performed for Gene Ontology (GO) terms—Molecular Function^96,97^, KEGG^98^, and Reactome^99^. The input for GO term enrichment consisted of Ensembl ID annotated ADO-specific DEGs tested against a background of 16,177 Ensembl-annotated genes. For KEGG and Reactome enrichment, which need Entrez gene IDs, the input comprised 360 ADO-specific DEGs (Entrez-annotated) against a background of 13,280 Entrez-annotated genes. This reduction in gene numbers for Entrez-based analyses was due to some Ensembl IDs lacking a direct Entrez ID mapping. All gene annotations were derived using the org.Mm.eg.db R package. The hypergeometric test was used to calculate significance, and analyses were restricted to terms or pathways containing 10-500 genes.

#### 5.5.6 Protein–protein interaction network analysis

A protein-protein interaction (PPI) network was constructed to map functional associations among proteins encoded by the ADO-specific DEGs. Interaction data were sourced from the STRING database (v12.0; minimum confidence score > 0.4)^100^. The resulting network was then imported into Cytoscape^101^, where the top seven hub proteins were identified using the Maximal Neighborhood Component (MNC) algorithm of the cytoHubba plugin (v0.1)^102^.

#### 5.5.7 Topology analysis

R package SEMgraph^103^ was used for topological analysis of RNA-Seq data. The analysis is predicted on three key inputs: a gene interaction network, the processed gene expression dataset, and the sample group design (in this instance, ADO (2h) versus CONTROL).

Molecular networks were constructed using pathway information from the Reactome databases using the Graphite^104^ R package. Pathways from this database were merged to create a comprehensive network. To prepare the network for directional analysis, bidirectional edges were removed, and only nodes (vertices) with at least one remaining edge were retained. This resulted in initial graphs for Reactome with 5,575 edges and 152,052 nodes.

For this analysis, gene expression data (previously filtered for low read counts and technical replicates collapsed) were further processed. Batch effects were adjusted using ComBat-seq function. Gene identifiers were converted to Entrez IDs. Subsequently, expression values were rank-based inverse normal transformed using the huge.npn function from the R package huge^105^ to mitigate normality constraints. The resulting expression matrix for topology analysis comprised 13,025 genes for the CONTROL and ADO (2h)-treated conditions (n=7 biological replicates per condition). Edges in the merged network were weighted based on gene expression data. Pairwise Pearson correlations between connected genes were calculated across samples, transformed using Fisher’s *r*-to-*z* transformation, and corresponding *z*-scores (zsign) and *p*-values were derived to represent edge weights and significance.

A Steiner Tree (ST) algorithm, specifically a fast Kou’s algorithm implementation, was employed to extract relevant subnetworks from the weighted graph, using the zsign values as edge weights.^106,107^ The ST approach identifies a minimal subnetwork connecting a predefined set of “seed” genes. In our case, the seed genes for each pathway were those that were part of the pathway and specific to the ADO treatment.

Extracted subnetworks were evaluated using Structural Equation Modeling (SEM) via the SEMrun function in the SEMgraph^103^ R package to assess both perturbation effects and overall model fit. Perturbation effects, representing local fit, were considered significant when the *p*-value was less than 0.05. The combined effect of all nodes was considered significantly activated if *pNodeAct* < 0.05 and/or inhibited if *pNodeIhn* < 0.05. Global model fit was assessed using the standardized root mean square residual (srmr) and the deviance-to-degrees-of-freedom ratio (dev/df). A srmr value below 0.1 and a dev/df ratio less than 3 were considered indicative of an acceptable global fit. It is noteworthy that global and local fit indices are not necessarily dependent; thus, even when global fit is suboptimal, locally significant relationships may still be valid and informative.

The resulting SEM graphs were visualized using the igraph^108^ R package. Nodes were color-coded to represent regulatory direction and significance: red for significant activation, blue for significant repression, and white for non-significant expression. Edges were in black if the connection between nodes is significant, with arrow and tee sign indicating positive and negative regulation.

## Data Availability

The original data of the work will be uploaded to GEO upon acceptance.

## Supporting Information

Supplementary figures include Mast cell protease expression, Principal component analysis for all conditions, Volcano plots comparing all four treated groups to CONTROL, and Expression profiling of inflammatory mediators regulated by ADO. Supplementary tables provide DESeq2 normalized expression of calcium ion channels in CONTROL, Differential expression statistics of all ADO-Specific protein coding Genes, Summary of top six most abundant protein classes among upregulated ADO-specific genes, Summary of top two most abundant protein classes among downregulated ADO-specific genes, and Summary of genes involved in inflammatory mediator biosynthesis and immune receptors in mast cells (PDF).

## Author Information

### Corresponding Author

Marc Freichel - Institute of Pharmacology, Heidelberg University

Andreas Keller - Clinical Bioinformatics, Saarland University

### Authors

Qihua Liang - Institute of Pharmacology, Heidelberg University

Volodymyr Tsvilovskyy - Institute of Pharmacology, Heidelberg University

Anouar Belkacemi - Institute of Pharmacology, Heidelberg University

Merima Bukva - Institute of Pharmacology, Heidelberg University

Christin Richter - Institute of Pharmacology, Heidelberg University

Nicole Ludwig - Clinical Bioinformatics, Saarland University

### Author Contributions

Conceptualization MF. Data Curation: QL, VT, MB, NL. Formal Analysis: QL, VT, MB, CR. Funding acquisition: AK, MF. Investigation: QL, VT, MB. Methodology: QL, VT, AB, CR, MB, NL. Supervision: VT, AB, AK, MF. Visualization: QL, VT, MF. Writing, original draft: QL, VT. Writing, review, and editing: QL, VT, AB, AK, MF.

### Notes

The authors declare no competing financial interest.

## ACKNOWLEDGMENT

We are thankful to Xenia Tolksdorf for technical assistance in mouse genotyping, PMCs isolation and cultivation as well as RNA isolation and cDNA libraries preparation. We are grateful to Vladimir Kuryshev for bioinformatical analysis of results of first test samples. We are thankful to EMBL Heidelberg Genomics Core Facility team and its Head Vladimir Benes for cDNA libraries sequencing and the team from the Interfakultäre Biomedizinische Forschungseinrichtung (IBF) from the Heidelberg University for expert technical assistance.

## ABBREVIATIONS

ADO: adenosine
BH: Benjamini-Hochberg
C48/80: Compound 48/80
DAG: diacylglycerol
DEGs: Differentially Expressed Genes
DNP: 2,4-dinitrophenyl human serum albumin
ER: endoplasmic reticulum
PLC: phospholipase C
PMCs: peritoneal mast cells
PMD: piecemeal degranulation
PIP2: phosphatidylinositol 4,5-bisphosphate
IgE: immunoglobulin E
IP3: inositol 1,4,5-trisphosphate
SOCE: store-operated calcium entry
TF: Transcription factor.

