## Supplementary for "Adenosine-specific transcriptional programs in murine connective tissue type mast cells"

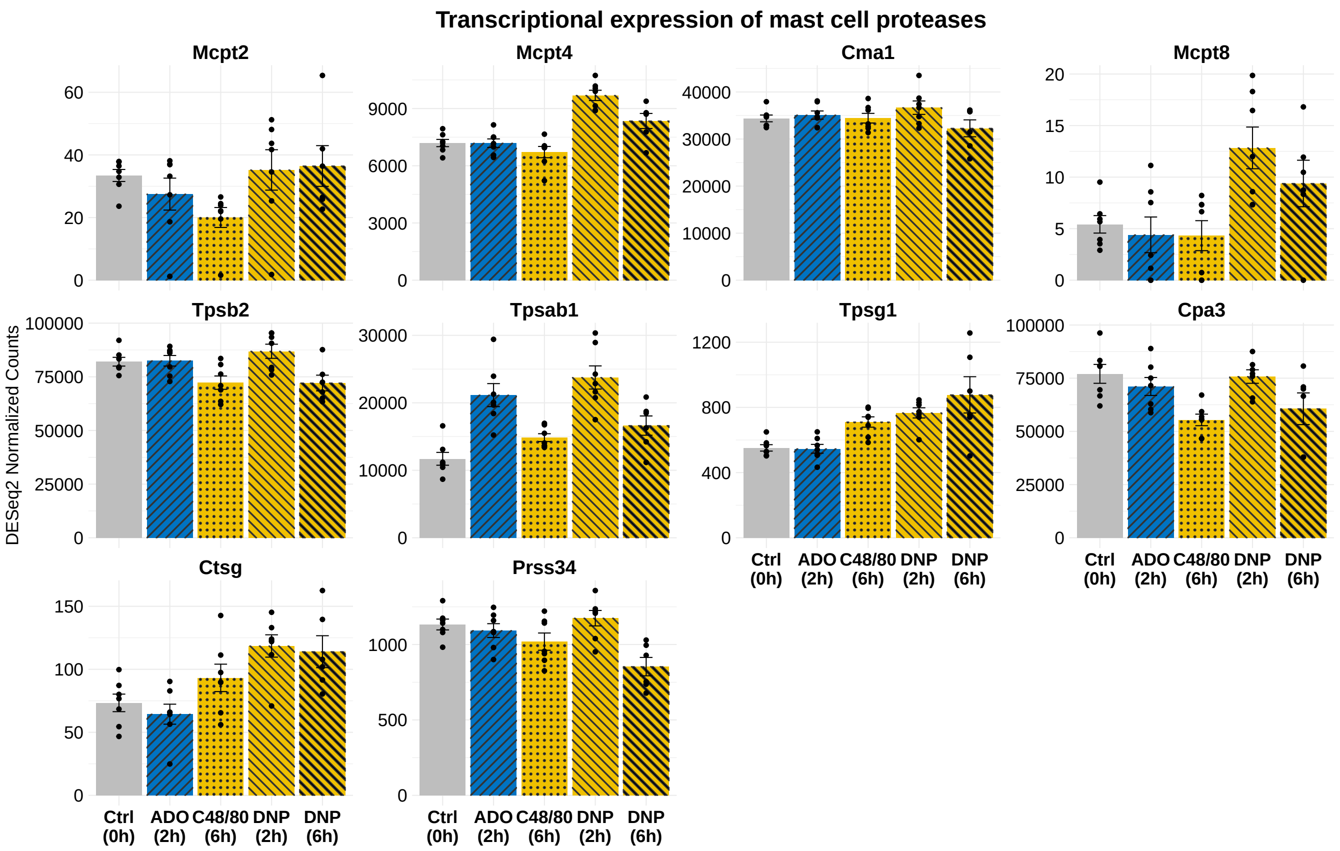


**Figure S1: Expression profiling of mast cell proteases in PMCs.**

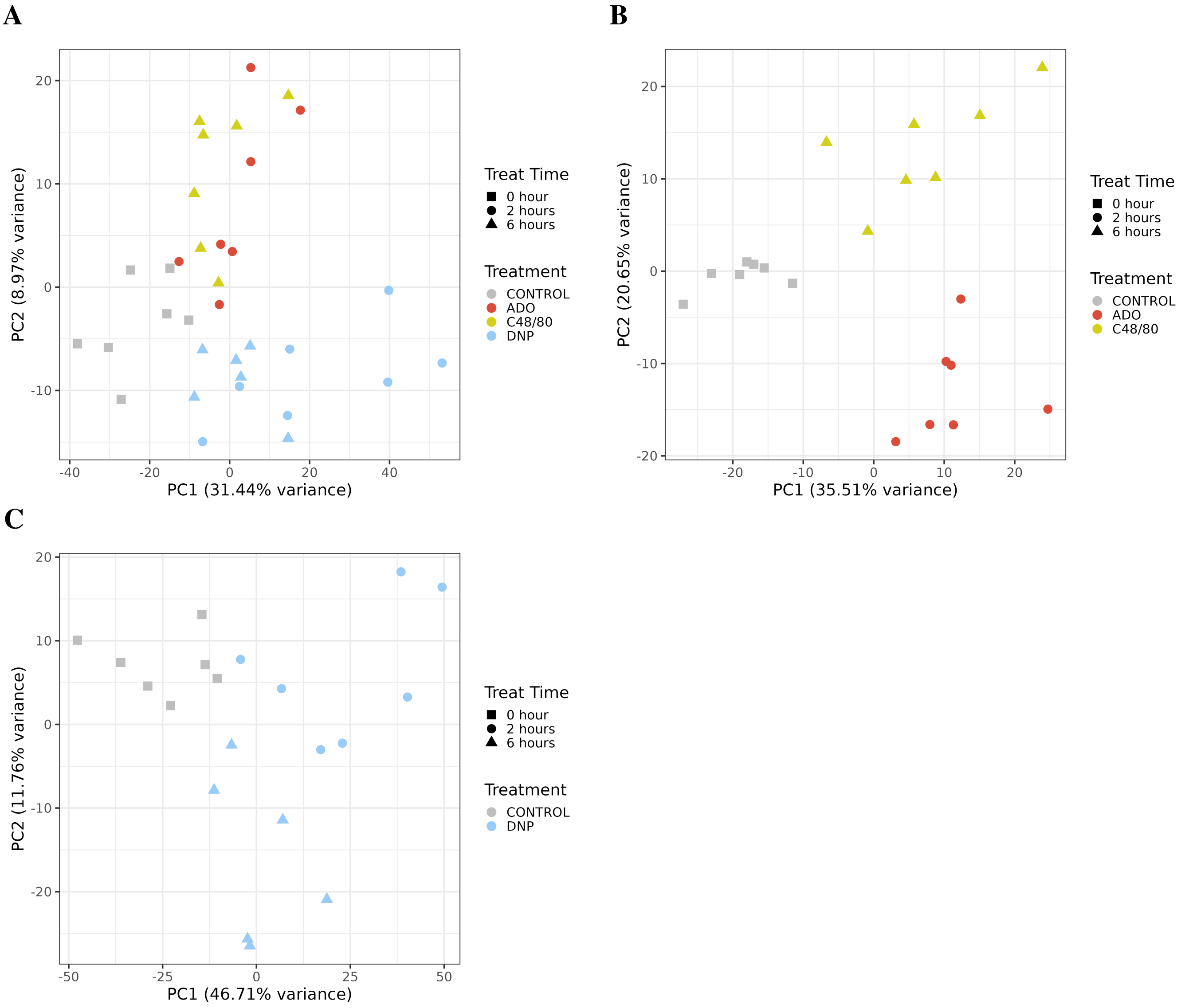


**Figure S2:** Principal component analysis of transcriptomic profiles following treatment with ADO, C48/80, and DNP. **A** PCA of all five experimental groups: CONTROL, ADO (2h), C48/80 (6h), DNP (2h), and DNP (6h). (B) PCA of CONTROL, ADO (2h), and C48/80 (6h) groups. (C) PCA of CONTROL, DNP (2h), and DNP (6h) groups. Each point represents an individual sample, colored by treatment and shaped according to treatment duration (square = 0 h, circle = 2 h, triangle = 6 h). Percent variance explained by each principal component is indicated on the axes.

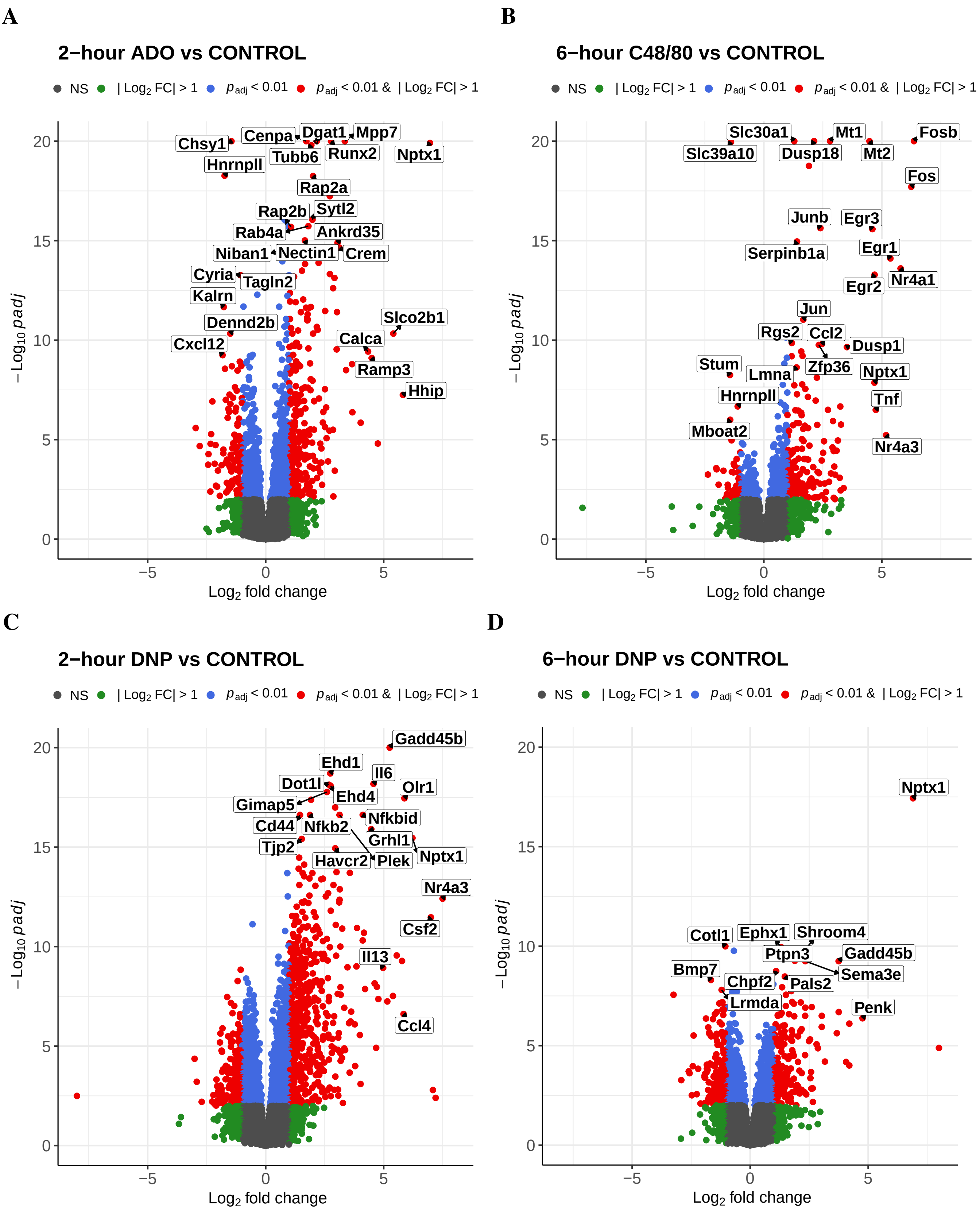


**Figure S3: Transcriptomic analysis reveals DEGs in stimulated mast cells**
Volcano plot of DEGs in mast cells following (**A**) 2-hour ADO treatment versus CONTROL. (**B**) 6-hour C48/80 treatment versus CONTROL. (**C**) 2-hour DNP treatment versus CONTROL. (**D**) 6-hour DNP treatment versus CONTROL. Significant DEGs fulfills $|\log_2 \mathrm{FC}| > 1$ and $p_{\mathrm{adj}} < 0.01$.

**A**

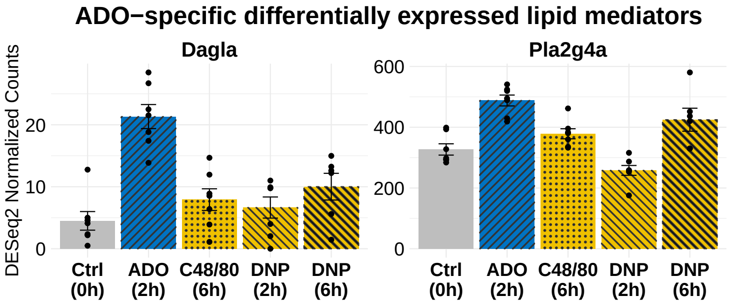
ADO−specific differentially expressed lipid mediators
Dagla
Pla2g4a
DESeq2 Normalized Counts
Ctrl (0h)
ADO (2h)
C48/80 (6h)
DNP (2h)
DNP (6h)

**B**

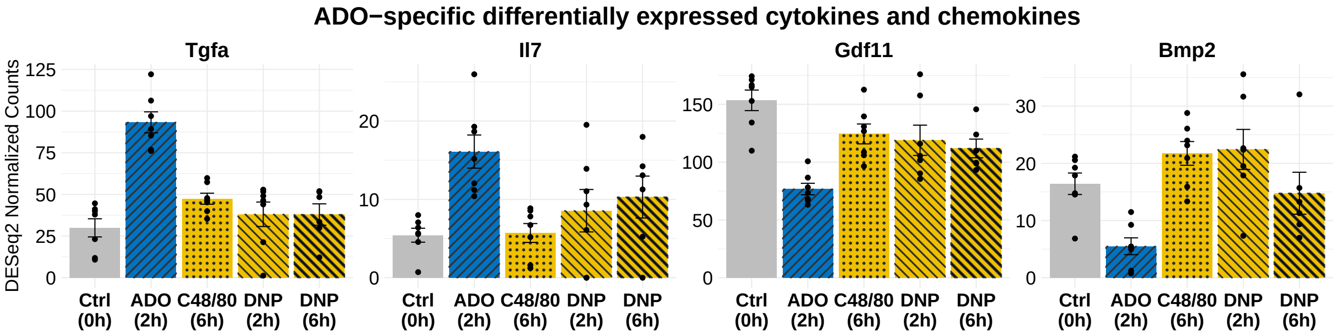
ADO−specific differentially expressed cytokines and chemokines
Tgfa
Il7
Gdf11
Bmp2
DESeq2 Normalized Counts
Ctrl (0h)
ADO (2h)
C48/80 (6h)
DNP (2h)
DNP (6h)

**C**

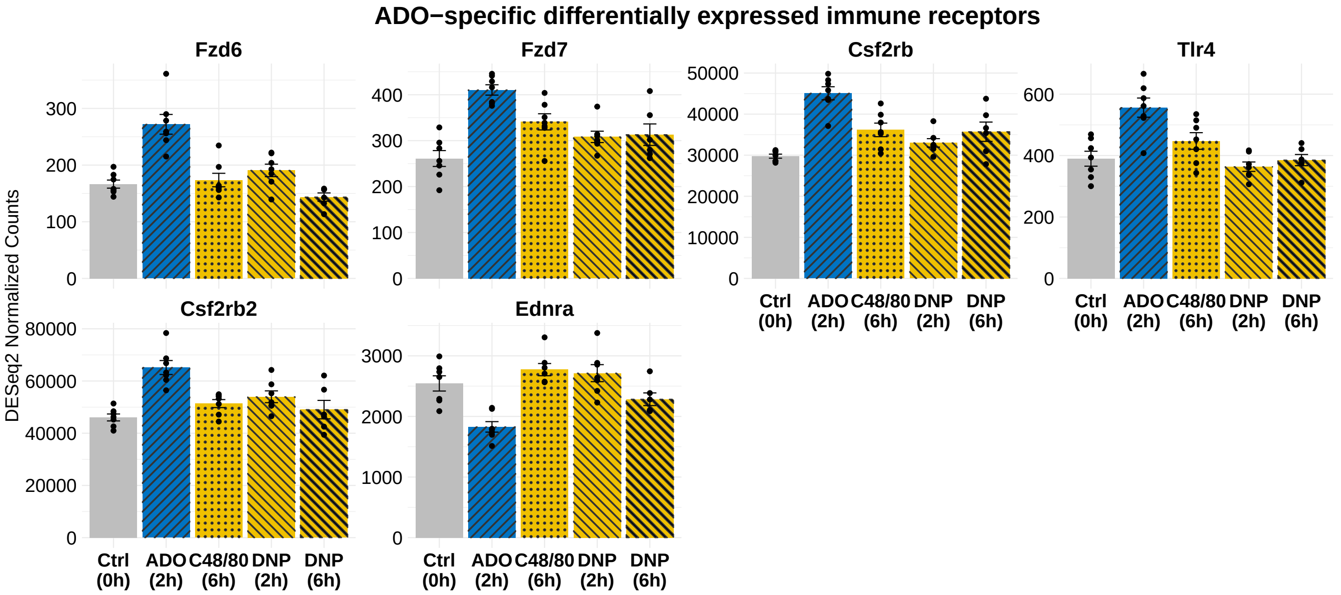
ADO−specific differentially expressed immune receptors
Fzd6
Fzd7
Csf2rb
Tlr4
Csf2rb2
Ednra
DESeq2 Normalized Counts
Ctrl (0h)
ADO (2h)
C48/80 (6h)
DNP (2h)
DNP (6h)

**D**

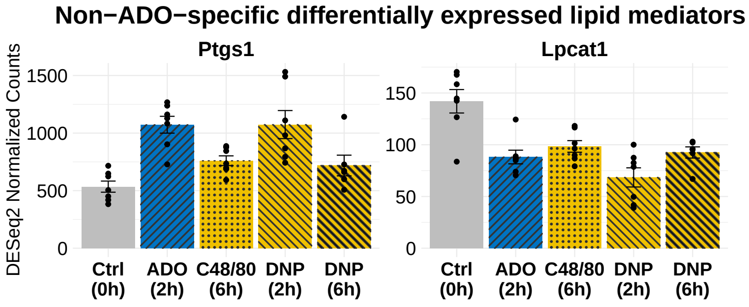
Non−ADO−specific differentially expressed lipid mediators
Ptgs1
Lpcat1
DESeq2 Normalized Counts
0
500
1000
1500
0
50
100
150
Ctrl (0h)
ADO (2h)
C48/80 (6h)
DNP (2h)
DNP (6h)

**E**

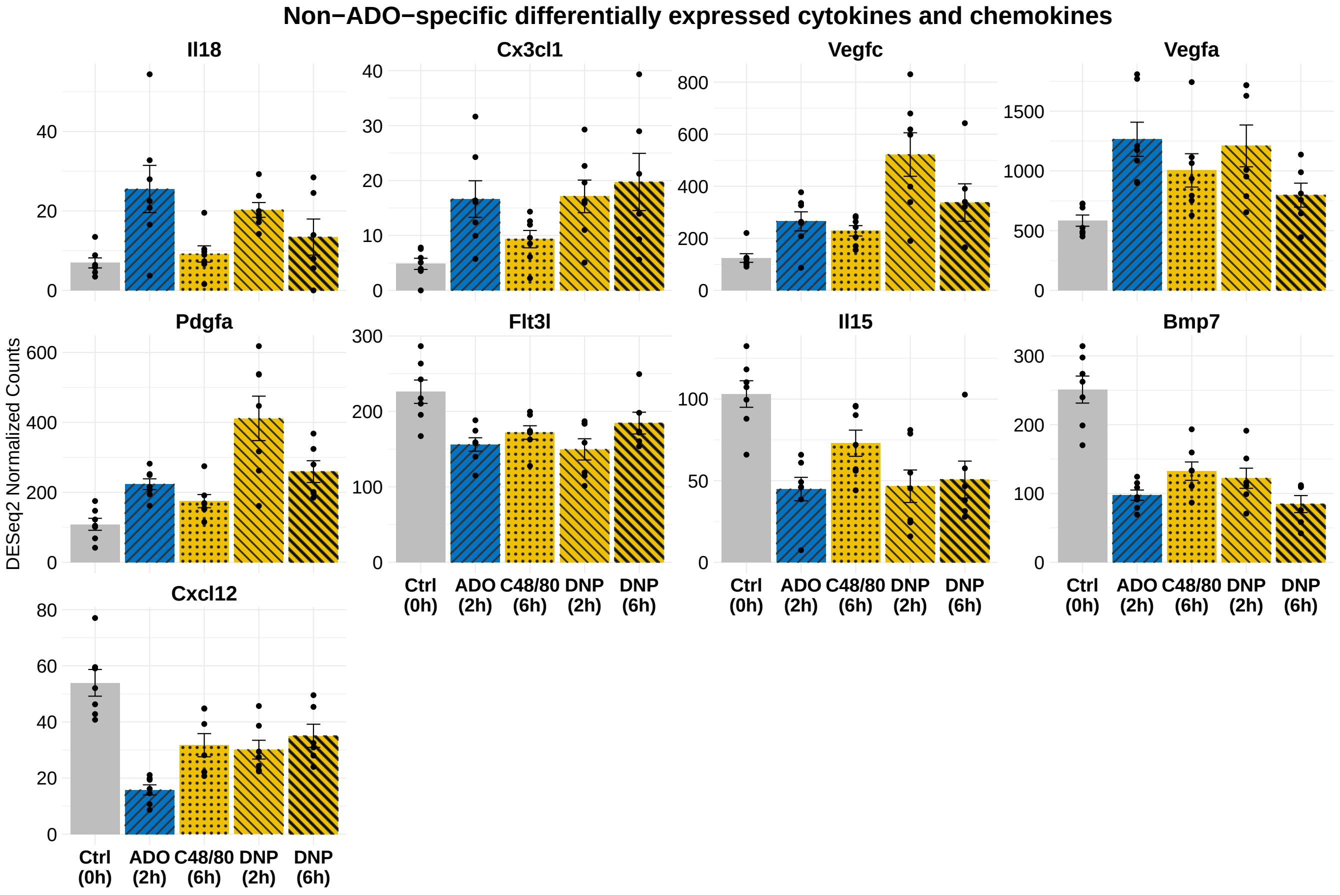
Non−ADO−specific differentially expressed cytokines and chemokines
Il18
Cx3cl1
Vegfc
Vegfa
Pdgfa
Flt3l
Il15
Bmp7
Cxcl12
DESeq2 Normalized Counts
Ctrl (0h)
ADO (2h)
C48/80 (6h)
DNP (2h)
DNP (6h)

**F**

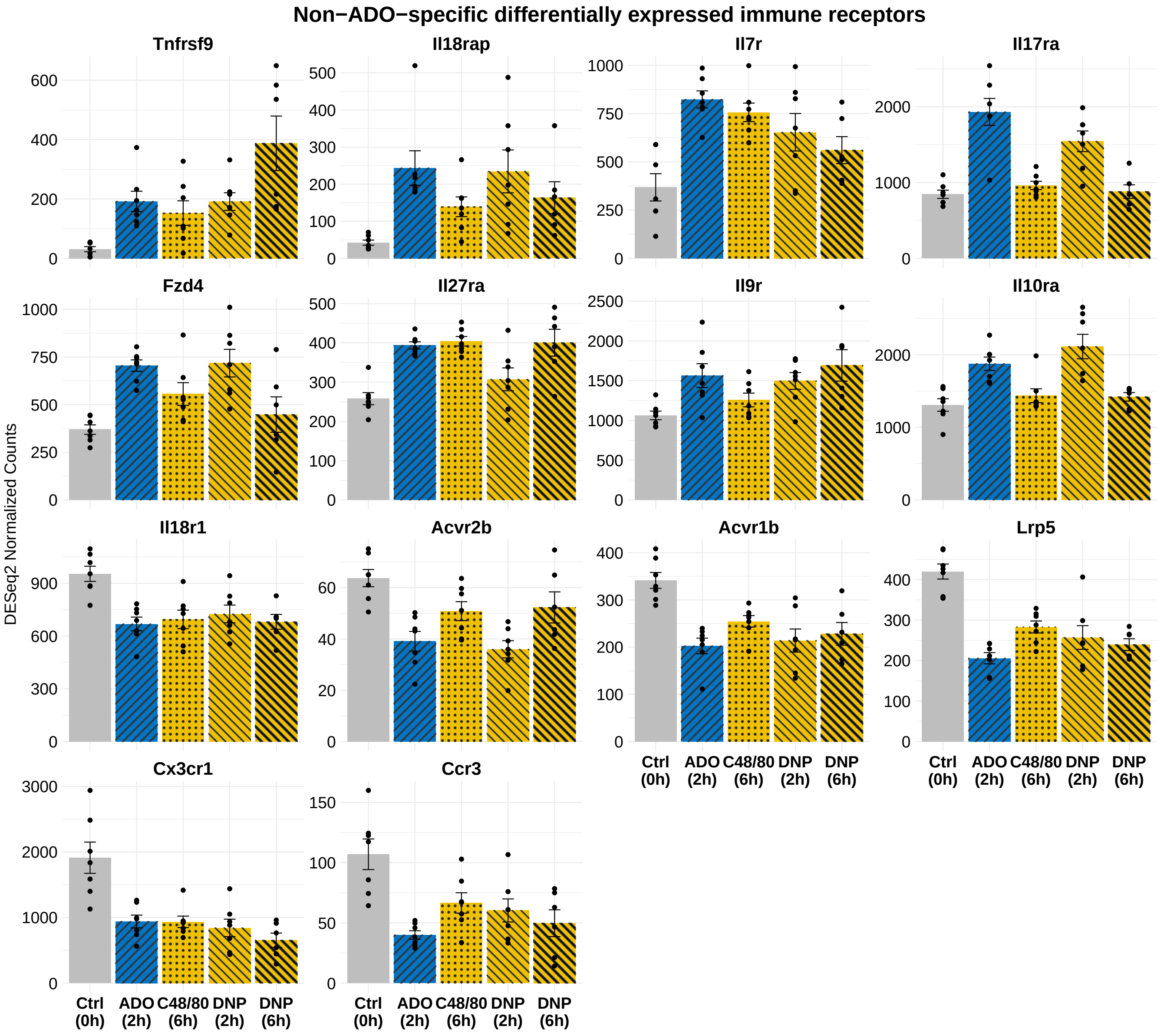


**Figure S4: Expression profiling of inflammatory mediators regulated by ADO in PMCs.**
**(A)** Lipid mediators significantly and exclusively regulated by ADO. **(B)** Cytokines, chemokines, or growth factors significantly and exclusively regulated by ADO. **(C)** Cytokine or chemokine receptors significantly and exclusively regulated by ADO. **(D)** Lipid mediators significantly, but not exclusively, regulated by ADO. **(E)** Cytokines, chemokines, or growth factors significantly, but not exclusively, regulated by ADO. **(F)** Cytokine or chemokine receptors significantly, but not exclusively, regulated by ADO. All significant regulation is defined as $p_{\text{adj}} < 0.01$. Genes were sorted in descending order according to $\log_2$ FC.

**Table S1:** DESeq2 normalized expression of calcium ion channels in CONTROL, ADO (2h), C48/80 (6h), DNP (2h), and DNP (6h)-treated samples. Genes that are differentially expressed compared to the CONTROL are marked with asterisks: * for $p_{\text{adj}} < 0.05$, ** for $p_{\text{adj}} < 0.01$, and *** for $p_{\text{adj}} < 0.001$.

| Gene Name | CONTROL | ADO (2h) | C48/80 (6h) | DNP (2h) | DNP (6h) |
|---|---|---|---|---|---|
| **Voltage-gated calcium channel $\alpha 1$ subunits (CACNA1 family)** | | | | | |
| Cacna1s | 1.0 | 1.5 | 1.9 | 1.3 | 1.8 |
| Cacna1c | 7.0 | 7.2 | 6.9 | 6.5 | 7.8 |
| Cacna1d | 41.9 | 54.2 | 43.2 | 39.1 | 36.1 |
| Cacna1f | 1.5 | 0.3 | 1.9 | 0.9 | 0.4 |
| Cacna1a | 16.7 | 13.8 | 14.8 | 11.4 | 13 |
| Cacna1b | 3.2 | 2.6 | 2.3 | 1.6 | 2.1 |
| Cacna1e | 54.1 | 54.6 | 52.4 | 33.6 | 33 |
| Cacna1g | 0.5 | 0.9 | 0.4 | 0 | 0 |
| Cacna1h | 54.5 | 72.9 | 82.4 | 108.3 | 103.7 |
| Cacna1i | 25.4 | 18.5 | 22.1 | 18.4 | 15.2 |
| **Voltage-gated calcium channel auxiliary $\beta$ subunits (CACNB family)** | | | | | |
| Cacnb1 | 10.9 | 7.1 | 7.5 | 12 | 8.9 |
| Cacnb2 | 10.4 | 9.1 | 8.2 | 10.2 | 12.2 |
| Cacnb3 | 7.7 | 7.5 | 8.4 | 8.8 | 9.8 |
| Cacnb4 | 261.7 | 263.4 | 188.2 * | 248 | 135.2 ** |
| **Voltage-gated calcium channel auxiliary $\alpha 2\delta$ subunits (CACNA2D family)** | | | | | |
| Cacna2d1 | 4.1 | 5.6 | 10.8 | 2.5 | 4.9 |
| Cacna2d2 | 0.6 | 0.7 | 0.7 | 2.7 | 0.9 |
| Cacna2d3 | 0.0 | 0 | 0 | 1 | 0.1 |
| Cacna2d4 | 16.8 | 10.9 | 13.5 | 16.3 | 10.1 |
| **Voltage-gated calcium channel auxiliary $\gamma$ subunits (CACNG family)** | | | | | |
| Cacng1 | 0.1 | 0.1 | 0 | 0 | 0.1 |
| Cacng2 | 0.0 | 0 | 0.3 | 0 | 0 |
| Cacng3 | 2.8 | 1.1 | 1.3 | 0.3 | 3.2 |
| Cacng4 | 0.1 | 0 | 0.2 | 0 | 0 |
| Cacng5 | 0.3 | 0 | 0.2 | 0 | 0.1 |
| Cacng6 | 0.6 | 0.1 | 0.5 | 0 | 0.3 |
| Cacng7 | 241.2 | 181.7 | 173.4 * | 200.8 | 137.2 *** |
| Cacng8 | 44.3 | 24.4 * | 25.8 | 26.5 * | 24.7 * |
| **Ryanodine receptors (RYR family)** | | | | | |
| Ryr1 | 2.0 | 1.5 | 3.1 | 1.2 | 0.9 |
| Ryr2 | 0.7 | 0.8 | 1.4 | 0.3 | 0.4 |
| Ryr3 | 6112.1 | 5642.1 | 9030.5 *** | 6649.1 | 9497.6 *** |
| **Inositol 1,4,5-trisphosphate receptors (ITPR family)** | | | | | |
| Itpr1 | 1683.7 | 1386.7 * | 1461.1 | 1608 | 1408.2 ** |
| Itpr2 | 4395.9 | 4008.7 | 4076.6 | 3894.7 | 3682.3 ** |
| Itpr3 | 547.7 | 551.3 | 652.4 | 540.6 | 791.7 ** |

Continued on next page

Table S1 – continued from previous page

| Gene Name | CONTROL | ADO (2h) | C48/80 (6h) | DNP (2h) | DNP (6h) |
|---|---|---|---|---|---|
| **Lysosomal calcium channels** | | | | | |
| Tpcn1 | 247.9 | 226.3 | 187.4 * | 204.6 | 206.9 |
| Tpcn2 | 122.8 | 114.9 | 120.8 | 114.6 | 112.9 |
| Tmem63a | 325.5 | 279.7 | 258.6 * | 269.4 | 257.6 * |
| Tmem63b | 141.5 | 199 * | 230.8 ** | 224.1 ** | 279.4 *** |
| **Transient receptor potential channels** | | | | | |
| Trpc1 | 0.8 | 2.2 | 1.3 | 2.2 | 1.1 |
| Trpc2 | 6.4 | 4.1 | 3.4 | 1.6 | 7.7 |
| Trpc3 | 0.3 | 0.1 | 0.2 | 0.4 | 0 |
| Trpc4 | 0.9 | 0.8 | 0.2 | 0.8 | 0.1 |
| Trpc5 | 4.9 | 2.4 | 4.5 | 2.5 | 2.8 |
| Trpc6 | 12.4 | 6.5 | 7.8 | 3.6 ** | 3.9 ** |
| Trpc7 | 45.2 | 30.9 | 23.7 | 31.6 | 15.7 ** |
| Trpv1 | 1.7 | 0.4 | 0.2 | 0.2 | 0.4 |
| Trpv2 | 212.3 | 245.4 | 260.1 | 265.9 | 166.1 |
| Trpv3 | 0.2 | 0 | 0 | 0 | 0.2 |
| Trpv4 | 2.4 | 3.8 | 3.7 | 1.1 | 1.2 |
| Trpv5 | 0.0 | 0 | 0 | 0 | 0 |
| Trpv6 | 0.4 | 0.4 | 0.1 | 0 | 0 |
| Trpm1 | 0.1 | 0.1 | 0.1 | 0 | 0 |
| Trpm2 | 113.4 | 83.7 | 65.6 * | 94.7 | 61.4 * |
| Trpm3 | 2.4 | 2.5 | 2.7 | 1.8 | 0.7 |
| Trpm4 | 267.6 | 459.7 ** | 252.1 | 498.9 *** | 375.3 |
| Trpm5 | 2.1 | 1.6 | 2.7 | 3.1 | 1.4 |
| Trpm6 | 0.0 | 0.2 | 0.2 | 0 | 0.1 |
| Trpm7 | 1721.6 | 1682.6 | 1682.3 | 1759.4 | 1815.9 |
| Trpm8 | 0.0 | 0 | 0.1 | 0 | 0 |
| Trpa1 | 0.0 | 0 | 0.2 | 0 | 0 |
| Mcoln1 | 377.0 | 303.5 * | 335 | 322.7 | 389.4 |
| Mcoln2 | 33.2 | 30.5 | 35.3 | 25.9 | 35.2 |
| Mcoln3 | 1.3 | 0.5 | 0.8 | 1.5 | 2.5 |
| Pkd2 | 412.1 | 376.4 | 364 | 378.5 | 368.8 |
| Pkd2l1 | 0.0 | 0 | 0 | 0 | 0 |
| Pkd2l2 | 65.7 | 58.2 | 42.2 | 62.6 | 49.9 |
| **Store-Operated Calcium Entry (SOCE)** | | | | | |
| Orai1 | 114.3 | 161.4 | 142.2 | 223.7 *** | 125.8 |
| Orai2 | 576.5 | 494.8 | 552 | 519.2 | 503.1 |
| Orai3 | 247.5 | 227.9 | 242.3 | 194.4 ** | 209.5 |
| Stim1 | 1526.0 | 1048.1 *** | 1295 | 1180.8 ** | 1101.2 *** |
| Stim2 | 491.7 | 502.9 | 487 | 703.9 *** | 522.9 |

Continued on next page

Table S1 – continued from previous page

| Gene Name | CONTROL | ADO (2h) | C48/80 (6h) | DNP (2h) | DNP (6h) |
|---|---|---|---|---|---|
| **P2X receptors (ATP-gated)** | | | | | |
| P2rx1 | 589.1 | 569.8 | 576.5 | 752.8 *** | 534.9 |
| P2rx2 | 0.3 | 0.2 | 0.3 | 0.2 | 0.3 |
| P2rx3 | 0.8 | 1.9 | 1.6 | 1.3 | 1.4 |
| P2rx4 | 2611.5 | 2091.5 * | 1879.2 ** | 1959.5 ** | 1994 * |
| P2rx5 | 7.8 | 10.5 | 9.2 | 13.4 | 8.5 |
| P2rx6 | 4.0 | 4.6 | 3.6 | 1.5 | 2.1 |
| P2rx7 | 2596.6 | 1937.1 * | 1966.2 * | 2117.6 * | 1974.7 * |
| **Piezo family** | | | | | |
| Piezo1 | 230.4 | 371.9 | 346.6 | 1646.3 *** | 506.7 * |
| Piezo2 | 6.1 | 6.4 | 29.6 | 9.6 | 6.7 |
| **NMDA receptors** | | | | | |
| Grin1 | 0.0 | 0.5 | 0 | 0 | 0 |
| Grin2a | 0.0 | 0.4 | 0 | 0 | 0 |
| Grin2b | 0.0 | 0 | 0 | 0 | 0.2 |
| Grin2c | 307.0 | 200.6 *** | 164.4 *** | 277.5 | 191.2 ** |
| Grin2d | 539.3 | 274.6 *** | 383.2 | 469.3 | 420.2 * |
| Grin3a | 26.0 | 23.7 | 22.5 | 24.8 | 22.7 |
| Grin3b | 3.7 | 3.4 | 3.5 | 4.8 | 2.9 |
| **AMPA receptor channel subunits** | | | | | |
| Gria1 | 1.5 | 1 | 1.3 | 1.1 | 3 |
| Gria2 | 0.3 | 0 | 0 | 0.2 | 0 |
| Gria3 | 19.8 | 20.4 | 33.2 | 23.4 | 22 |
| Gria4 | 0.3 | 0.1 | 0.3 | 0 | 0.5 |
| **Calcium homeostasis modulator** | | | | | |
| Calhm1 | 0.1 | 0.1 | 0 | 0.3 | 0 |
| Calhm2 | 526.7 | 332.9 ** | 381.9 | 361.5 ** | 393.9 |
| Calhm3 | 0.0 | 0 | 0 | 0 | 0 |
| Calhm4 | 0.0 | 0 | 0 | 0 | 0.1 |
| Calhm5 | 0.1 | 0.5 | 0.1 | 0 | 0 |
| Calhm6 | 0.4 | 0.9 | 0.5 | 0.4 | 0.7 |

**Table S2:** Differential expression statistics of all ADO-specific protein coding genes. Genes were ordered in terms of their normalized expression value in either CONTROL or ADO 2 hour-treated condition.

| Gene Name | CONTROL | ADO (2h) | log2FC | padj |
|---|---|---|---|---|
| Csf2rb2 | 46038.7 | 65174.7 | 0.5 | 2.0e-05 |
| Csf2rb | 29766.2 | 45078.7 | 0.6 | 3.5e-08 |
| Gata2 | 14346.1 | 10470.5 | -0.5 | 1.4e-05 |
| Bod1l | 5279.0 | 9133.9 | 0.8 | 7.1e-05 |
| Ywhaz | 6013.5 | 8442.7 | 0.5 | 3.2e-05 |
| Mtss1 | 7503.2 | 5092.1 | -0.6 | 5.5e-10 |
| Aldoa | 5709.2 | 7296.7 | 0.4 | 4.1e-03 |
| Bltp1 | 4509.3 | 7196.8 | 0.7 | 3.7e-06 |
| Notch2 | 5695.8 | 7134.9 | 0.3 | 7.8e-03 |
| Rab27b | 4326.2 | 6951.7 | 0.7 | 7.6e-06 |
| Niban1 | 2912.3 | 6381.1 | 1.1 | 2.3e-15 |
| Zbtb38 | 3234.7 | 5941.0 | 0.9 | 1.9e-16 |
| Prkar1a | 3784.5 | 4831.1 | 0.3 | 3.2e-03 |
| Ikzf2 | 4658.4 | 3452.4 | -0.4 | 7.4e-03 |
| Ldha | 2525.0 | 4618.7 | 0.8 | 9.8e-11 |
| Pfkp | 2521.4 | 4204.5 | 0.7 | 8.1e-08 |
| Ubash3b | 4110.8 | 2851.0 | -0.5 | 3.5e-08 |
| Paxbp1 | 2472.8 | 4110.2 | 0.7 | 9.7e-05 |
| Sdcbp | 2862.8 | 3910.0 | 0.5 | 3.8e-04 |
| Trio | 2338.3 | 3815.6 | 0.7 | 5.4e-06 |
| Rfx7 | 3637.1 | 2698.2 | -0.4 | 2.7e-06 |
| Itch | 3491.3 | 2557.6 | -0.4 | 3.1e-07 |
| Fryl | 3477.7 | 2860.3 | -0.3 | 2.3e-04 |
| Arrb1 | 2578.0 | 3396.6 | 0.4 | 5.9e-06 |
| Otulinl | 2299.3 | 3383.4 | 0.5 | 3.7e-04 |
| Ston2 | 1846.1 | 3371.9 | 0.8 | 3.7e-04 |
| Gvin1 | 1830.2 | 3052.5 | 0.7 | 5.3e-03 |
| Tnpo1 | 2405.6 | 2823.8 | 0.2 | 8.7e-03 |
| Prkcd | 2804.9 | 2089.5 | -0.4 | 2.8e-04 |
| Vat1 | 2096.1 | 2785.4 | 0.4 | 3.2e-03 |
| Taok3 | 2759.7 | 2317.6 | -0.3 | 6.6e-04 |
| Gvin2 | 1314.7 | 2701.0 | 1.0 | 2.8e-03 |
| Myo5a | 2190.1 | 2615.1 | 0.3 | 8.0e-03 |
| Ptpn13 | 1260.8 | 2589.0 | 1.0 | 4.7e-09 |
| Ednra | 2544.4 | 1827.7 | -0.5 | 9.1e-05 |
| Mrgprx2 | 1619.1 | 2460.4 | 0.6 | 3.7e-03 |
| Sema4d | 2452.8 | 1673.0 | -0.5 | 3.3e-04 |
| Arid4b | 2354.4 | 1794.7 | -0.4 | 2.0e-04 |
| Plcg1 | 2337.5 | 1644.4 | -0.5 | 1.0e-06 |

Continued on next page

Table S2 – continued from previous page

| Gene Name | CONTROL | ADO (2h) | log2FC | padj |
|---|---|---|---|---|
| Mat2a | 2322.7 | 1746.7 | -0.4 | 8.9e-03 |
| Gab2 | 1651.4 | 2322.5 | 0.5 | 4.5e-03 |
| Top1 | 2239.5 | 1618.0 | -0.5 | 2.1e-03 |
| Tns3 | 2140.6 | 1512.9 | -0.5 | 3.2e-05 |
| Arap1 | 2009.3 | 1420.0 | -0.5 | 8.1e-07 |
| Sipa1l1 | 1113.4 | 1936.3 | 0.8 | 6.1e-05 |
| Gmfb | 1603.4 | 1918.7 | 0.3 | 5.6e-03 |
| Nckap1 | 1862.9 | 1456.6 | -0.4 | 5.2e-13 |
| Cic | 1819.1 | 1458.2 | -0.3 | 5.8e-04 |
| Hs6st1 | 1055.1 | 1783.7 | 0.7 | 3.3e-04 |
| Ap1g1 | 1380.7 | 1682.1 | 0.3 | 9.0e-04 |
| Nup153 | 1678.0 | 1368.7 | -0.3 | 2.8e-03 |
| Mau2 | 1352.9 | 1669.4 | 0.3 | 4.9e-04 |
| Fermt3 | 1631.7 | 1322.0 | -0.3 | 3.5e-06 |
| Tiparp | 1592.6 | 978.6 | -0.7 | 4.6e-04 |
| Serpinb6a | 1126.3 | 1573.2 | 0.5 | 2.3e-04 |
| Nrdc | 1378.3 | 1571.9 | 0.2 | 8.1e-03 |
| Acap2 | 1460.6 | 1203.2 | -0.3 | 1.9e-03 |
| Ccser2 | 1154.4 | 1449.5 | 0.3 | 8.3e-04 |
| Coro1c | 1125.9 | 1443.3 | 0.4 | 2.3e-03 |
| P4ha1 | 1412.2 | 1142.9 | -0.3 | 1.3e-05 |
| Arpc3 | 905.2 | 1403.6 | 0.6 | 1.8e-06 |
| Isy1 | 284.8 | 1371.3 | 2.2 | 1.3e-14 |
| Arhgap25 | 1052.2 | 1355.5 | 0.4 | 1.5e-03 |
| Srpk1 | 1349.8 | 1151.0 | -0.2 | 3.1e-03 |
| Phldb3 | 891.9 | 1338.7 | 0.6 | 2.5e-05 |
| Nceh1 | 946.7 | 1303.3 | 0.5 | 3.6e-03 |
| Kif13b | 1291.2 | 965.7 | -0.4 | 2.8e-03 |
| Gga2 | 695.6 | 1272.8 | 0.9 | 9.2e-08 |
| Crem | 142.8 | 1265.5 | 3.1 | 2.3e-15 |
| Tmed5 | 869.9 | 1248.7 | 0.5 | 6.2e-03 |
| Lxn | 767.1 | 1244.4 | 0.7 | 2.8e-07 |
| Dstyk | 1033.3 | 1232.7 | 0.3 | 5.7e-03 |
| Coro2a | 1216.7 | 702.6 | -0.7 | 6.1e-03 |
| Itk | 495.5 | 1198.1 | 1.2 | 6.3e-04 |
| Gak | 979.7 | 1191.1 | 0.3 | 4.0e-03 |
| Pou2f1 | 896.0 | 1187.8 | 0.4 | 4.1e-03 |
| Plekha2 | 569.0 | 1180.3 | 1.1 | 2.0e-09 |
| Slc2a1 | 1162.1 | 808.9 | -0.5 | 2.3e-03 |
| Dctn4 | 881.9 | 1145.2 | 0.4 | 2.0e-04 |
| Mbtps1 | 1131.0 | 980.4 | -0.2 | 9.1e-03 |

Continued on next page

Table S2 – continued from previous page

| Gene Name | CONTROL | ADO (2h) | log2FC | padj |
|---|---|---|---|---|
| Map2k4 | 937.6 | 1122.9 | 0.3 | 9.5e-03 |
| Gpbp1l1 | 1096.3 | 938.6 | -0.2 | 8.9e-03 |
| Nedd4l | 704.3 | 1014.6 | 0.5 | 4.1e-03 |
| Sbno2 | 748.1 | 1012.3 | 0.5 | 1.1e-05 |
| Slc25a5 | 644.7 | 1004.5 | 0.6 | 2.7e-04 |
| Stat4 | 656.7 | 1001.8 | 0.6 | 2.0e-03 |
| St3gal2 | 989.3 | 667.0 | -0.6 | 3.0e-03 |
| Rap2b | 466.8 | 972.7 | 1.1 | 2.0e-16 |
| Cd300lb | 456.7 | 946.3 | 1.1 | 1.0e-07 |
| Srek1 | 943.5 | 758.7 | -0.3 | 7.8e-03 |
| Lrrc1 | 619.9 | 943.0 | 0.6 | 1.3e-05 |
| Ttpal | 671.6 | 936.7 | 0.5 | 2.6e-03 |
| Vamp4 | 632.0 | 930.4 | 0.6 | 1.5e-10 |
| Flcn | 922.0 | 768.0 | -0.3 | 7.9e-03 |
| Pmp22 | 485.8 | 918.2 | 0.9 | 2.8e-07 |
| Zfp398 | 764.8 | 906.9 | 0.3 | 3.8e-03 |
| Kpna1 | 618.3 | 880.4 | 0.5 | 5.8e-04 |
| Osbpl9 | 580.5 | 873.2 | 0.6 | 1.3e-05 |
| Jade1 | 643.2 | 872.9 | 0.4 | 2.6e-04 |
| Wdr81 | 872.5 | 672.1 | -0.4 | 9.6e-04 |
| Elovl5 | 662.2 | 865.2 | 0.4 | 5.2e-03 |
| Ntng2 | 396.7 | 851.1 | 1.1 | 2.8e-04 |
| Wbp2 | 846.4 | 636.9 | -0.4 | 1.7e-03 |
| Klf10 | 478.7 | 841.4 | 0.8 | 2.2e-06 |
| Med14 | 605.5 | 830.1 | 0.4 | 2.9e-03 |
| Kdm4b | 827.9 | 637.1 | -0.4 | 8.5e-03 |
| Nap1l5 | 219.3 | 821.2 | 1.9 | 1.5e-09 |
| Tpm4 | 335.6 | 811.8 | 1.2 | 1.0e-05 |
| Sntb2 | 578.4 | 802.5 | 0.5 | 1.1e-04 |
| Amph | 463.5 | 791.6 | 0.8 | 4.3e-03 |
| Mpp7 | 77.1 | 781.3 | 3.4 | 3.1e-38 |
| Cpt1a | 600.0 | 780.1 | 0.4 | 9.0e-04 |
| Slc36a4 | 661.5 | 777.4 | 0.2 | 5.6e-03 |
| Scin | 476.3 | 763.2 | 0.7 | 5.2e-03 |
| Dhrs3 | 760.0 | 445.4 | -0.8 | 6.9e-03 |
| Card11 | 747.7 | 389.0 | -0.9 | 2.3e-08 |
| Pitpnc1 | 565.8 | 747.6 | 0.4 | 7.5e-03 |
| Galc | 415.1 | 727.3 | 0.8 | 2.9e-06 |
| Plin2 | 415.9 | 722.9 | 0.8 | 8.9e-05 |
| Rrp1 | 568.6 | 719.7 | 0.3 | 4.7e-03 |
| Pik3cb | 524.8 | 713.9 | 0.4 | 9.8e-04 |

Continued on next page

Table S2 – continued from previous page

| Gene Name | CONTROL | ADO (2h) | log2FC | padj |
|---|---|---|---|---|
| Ric1 | 709.8 | 548.8 | -0.4 | 5.4e-04 |
| Gusb | 474.4 | 701.6 | 0.5 | 2.2e-03 |
| Aida | 537.0 | 691.3 | 0.4 | 6.3e-04 |
| Sap130 | 690.1 | 560.5 | -0.3 | 3.5e-03 |
| Limd2 | 682.8 | 484.8 | -0.5 | 3.1e-05 |
| Qser1 | 543.4 | 680.0 | 0.3 | 9.8e-03 |
| Calcoco1 | 678.3 | 487.9 | -0.5 | 1.2e-03 |
| Rfk | 482.0 | 671.2 | 0.5 | 6.9e-04 |
| Ulk1 | 662.6 | 485.7 | -0.4 | 8.2e-03 |
| Rfx5 | 648.4 | 511.4 | -0.3 | 2.7e-04 |
| Pi4kb | 470.6 | 641.8 | 0.4 | 1.1e-03 |
| Mink1 | 635.9 | 430.1 | -0.6 | 3.5e-05 |
| Ctla2b | 354.1 | 622.4 | 0.8 | 1.3e-04 |
| Lrch4 | 612.0 | 375.2 | -0.7 | 2.7e-03 |
| Stx7 | 517.2 | 598.3 | 0.2 | 6.4e-03 |
| Vdac3 | 493.7 | 597.8 | 0.3 | 3.5e-03 |
| Pts | 465.8 | 593.5 | 0.4 | 3.2e-03 |
| Man1b1 | 443.9 | 588.7 | 0.4 | 1.3e-03 |
| Tbcel | 585.4 | 488.3 | -0.3 | 7.7e-03 |
| Disp1 | 361.8 | 574.2 | 0.7 | 2.4e-10 |
| Entpd7 | 355.6 | 559.4 | 0.6 | 3.8e-07 |
| Tlr4 | 389.8 | 556.2 | 0.5 | 5.2e-04 |
| Rnf217 | 355.8 | 535.5 | 0.6 | 1.4e-06 |
| Cemip2 | 520.8 | 321.7 | -0.7 | 5.9e-04 |
| Golt1b | 415.9 | 519.8 | 0.3 | 5.1e-03 |
| Antxr2 | 329.3 | 516.2 | 0.7 | 9.8e-03 |
| Strap | 374.6 | 512.3 | 0.5 | 2.9e-03 |
| Mcur1 | 501.2 | 411.4 | -0.3 | 3.5e-03 |
| Rassf3 | 345.8 | 496.9 | 0.5 | 4.7e-03 |
| Rhoq | 333.5 | 496.0 | 0.6 | 7.2e-03 |
| Grin2d | 495.1 | 244.8 | -1.0 | 9.8e-06 |
| Ninj1 | 261.5 | 493.8 | 0.9 | 1.6e-07 |
| Asb1 | 489.0 | 353.0 | -0.5 | 7.8e-04 |
| Unc45a | 378.0 | 488.8 | 0.4 | 1.5e-03 |
| Pla2g4a | 326.9 | 488.1 | 0.6 | 3.6e-04 |
| Emd | 245.6 | 487.5 | 1.0 | 5.5e-14 |
| Plekha5 | 297.3 | 481.2 | 0.7 | 2.5e-04 |
| Gucd1 | 365.5 | 479.1 | 0.4 | 5.2e-03 |
| Ksr1 | 474.0 | 291.8 | -0.7 | 1.8e-04 |
| Adam9 | 353.0 | 472.6 | 0.4 | 6.2e-03 |
| Hpcal1 | 142.8 | 470.0 | 1.8 | 7.6e-12 |

Continued on next page

Table S2 – continued from previous page

| Gene Name | CONTROL | ADO (2h) | log2FC | padj |
|---|---|---|---|---|
| Slc2a6 | 467.8 | 294.3 | -0.6 | 3.1e-06 |
| Tmem184c | 466.9 | 358.0 | -0.4 | 8.0e-03 |
| Pias2 | 466.0 | 398.2 | -0.2 | 7.8e-03 |
| Prune1 | 360.3 | 465.7 | 0.4 | 5.6e-03 |
| Atp5f1d | 327.7 | 456.1 | 0.5 | 2.1e-03 |
| Inip | 316.3 | 451.5 | 0.5 | 7.2e-04 |
| Wdr44 | 362.1 | 450.6 | 0.3 | 4.2e-03 |
| Smim30 | 358.0 | 450.0 | 0.3 | 2.0e-03 |
| Map4k2 | 449.5 | 327.9 | -0.5 | 7.1e-03 |
| Dgka | 449.3 | 330.1 | -0.5 | 4.0e-03 |
| Ptch1 | 435.2 | 308.0 | -0.5 | 1.0e-04 |
| Fbxo9 | 250.1 | 432.6 | 0.8 | 8.9e-17 |
| Gata1 | 431.4 | 247.4 | -0.8 | 9.9e-04 |
| Eif1ax | 326.7 | 425.5 | 0.4 | 1.8e-03 |
| Zeb1 | 423.7 | 321.5 | -0.4 | 2.2e-03 |
| Ncoa7 | 416.0 | 260.2 | -0.7 | 2.3e-09 |
| Nol4l | 415.2 | 257.9 | -0.7 | 1.1e-04 |
| Tbc1d31 | 287.9 | 414.5 | 0.5 | 4.7e-03 |
| Fzd7 | 261.0 | 410.4 | 0.6 | 6.7e-06 |
| Bcl9 | 405.9 | 250.0 | -0.7 | 7.2e-09 |
| Prep | 254.1 | 396.4 | 0.6 | 5.4e-04 |
| Txndc5 | 298.9 | 389.5 | 0.4 | 6.6e-03 |
| Lztfl1 | 311.0 | 388.0 | 0.3 | 7.6e-04 |
| Plcb3 | 387.2 | 254.6 | -0.6 | 1.7e-06 |
| Zcchc10 | 383.8 | 213.1 | -0.8 | 4.8e-05 |
| Nfix | 383.3 | 289.8 | -0.4 | 3.6e-03 |
| Pcgf5 | 249.5 | 370.4 | 0.6 | 4.6e-04 |
| Slc6a12 | 364.4 | 231.7 | -0.7 | 2.3e-03 |
| Tesk2 | 363.2 | 246.0 | -0.6 | 6.5e-04 |
| Ccnc | 275.9 | 359.6 | 0.4 | 2.8e-03 |
| Rbm45 | 212.1 | 350.8 | 0.7 | 1.6e-05 |
| Eeig1 | 350.2 | 212.4 | -0.7 | 6.3e-04 |
| Osbpl6 | 28.2 | 346.2 | 3.7 | 1.6e-09 |
| Apex2 | 340.9 | 244.1 | -0.5 | 5.5e-03 |
| Fcho1 | 336.7 | 275.2 | -0.3 | 8.0e-03 |
| Ptov1 | 335.0 | 257.4 | -0.4 | 6.7e-03 |
| Slc25a37 | 233.8 | 329.7 | 0.5 | 4.8e-04 |
| Tshz3 | 328.8 | 214.2 | -0.6 | 6.2e-06 |
| Impdh1 | 216.0 | 318.0 | 0.5 | 1.8e-04 |
| Bcorl1 | 315.8 | 206.4 | -0.6 | 7.0e-05 |
| Pgpep1 | 312.7 | 201.6 | -0.6 | 9.4e-05 |

Continued on next page

Table S2 – continued from previous page

| Gene Name | CONTROL | ADO (2h) | log2FC | padj |
|---|---|---|---|---|
| Mfhas1 | 208.8 | 312.7 | 0.6 | 2.7e-04 |
| Slc20a2 | 224.7 | 311.3 | 0.5 | 9.7e-03 |
| Osgin1 | 305.1 | 167.6 | -0.8 | 3.7e-03 |
| Sdhb | 226.2 | 303.5 | 0.4 | 2.0e-04 |
| Tinf2 | 206.5 | 290.2 | 0.5 | 3.4e-04 |
| Ccdc167 | 288.4 | 221.0 | -0.4 | 6.5e-03 |
| Rnf157 | 288.2 | 206.4 | -0.5 | 1.1e-03 |
| Bbs12 | 199.9 | 284.0 | 0.5 | 2.7e-03 |
| Cyth1 | 196.1 | 275.9 | 0.5 | 3.2e-03 |
| Ap5z1 | 273.7 | 206.7 | -0.4 | 9.0e-03 |
| Smg9 | 215.5 | 272.3 | 0.3 | 6.1e-03 |
| Fzd6 | 166.5 | 272.0 | 0.7 | 1.5e-06 |
| Nfkbie | 269.6 | 187.1 | -0.5 | 4.0e-03 |
| Aph1c | 158.9 | 268.2 | 0.8 | 2.4e-06 |
| Phlpp2 | 266.0 | 191.9 | -0.5 | 2.8e-03 |
| Tirap | 150.4 | 263.8 | 0.8 | 7.1e-09 |
| Hdac7 | 263.8 | 183.7 | -0.5 | 9.5e-04 |
| Zfp956 | 145.6 | 261.7 | 0.9 | 8.7e-12 |
| Nsun4 | 160.0 | 254.5 | 0.7 | 4.1e-07 |
| Rabep2 | 251.1 | 181.7 | -0.5 | 1.4e-03 |
| Unc119b | 198.2 | 249.3 | 0.3 | 1.2e-03 |
| Grm5 | 106.9 | 244.7 | 1.2 | 3.9e-03 |
| Tmc4 | 146.0 | 243.2 | 0.7 | 7.0e-03 |
| Crtc1 | 242.7 | 183.1 | -0.4 | 8.2e-03 |
| Lactb2 | 180.8 | 241.1 | 0.4 | 1.1e-03 |
| Plxnb3 | 239.6 | 125.0 | -1.0 | 3.0e-03 |
| Zfhx2 | 238.7 | 165.3 | -0.6 | 4.6e-03 |
| Lrrc57 | 238.4 | 160.3 | -0.6 | 1.7e-06 |
| Card19 | 158.8 | 238.3 | 0.6 | 6.6e-03 |
| Ahdc1 | 236.0 | 141.7 | -0.7 | 5.5e-03 |
| Mfsd12 | 119.8 | 232.3 | 1.0 | 7.6e-08 |
| Tpbgl | 106.5 | 231.5 | 1.1 | 7.0e-04 |
| Dlg4 | 231.3 | 159.1 | -0.5 | 7.8e-05 |
| Arf2 | 143.4 | 230.8 | 0.7 | 7.3e-04 |
| Lmnb1 | 82.9 | 216.8 | 1.3 | 3.0e-05 |
| Kcnn4 | 135.2 | 216.7 | 0.7 | 1.1e-03 |
| Gcsh | 74.3 | 208.7 | 1.5 | 3.9e-12 |
| Stk39 | 208.3 | 151.2 | -0.4 | 2.8e-03 |
| Slc16a3 | 143.6 | 207.8 | 0.5 | 1.4e-03 |
| Mpped2 | 204.0 | 150.2 | -0.4 | 5.3e-03 |
| Wdr90 | 99.2 | 201.2 | 1.0 | 1.9e-05 |

Continued on next page

Table S2 – continued from previous page

| Gene Name | CONTROL | ADO (2h) | log2FC | padj |
|---|---|---|---|---|
| Zcchc14 | 154.8 | 200.8 | 0.4 | 2.4e-03 |
| Dgke | 147.8 | 197.3 | 0.4 | 8.1e-03 |
| Rab4a | 56.3 | 194.3 | 1.8 | 1.9e-16 |
| Usp20 | 191.9 | 134.7 | -0.5 | 2.3e-03 |
| Zfp619 | 191.2 | 139.0 | -0.5 | 7.7e-03 |
| Klhl23 | 132.0 | 189.6 | 0.5 | 2.3e-03 |
| Aatk | 187.5 | 69.4 | -1.4 | 2.7e-05 |
| Axin2 | 187.4 | 130.6 | -0.5 | 6.7e-03 |
| Pnkd | 180.3 | 107.5 | -0.7 | 1.7e-03 |
| Insig1 | 178.8 | 103.3 | -0.8 | 1.3e-03 |
| Ifih1 | 126.0 | 178.5 | 0.5 | 1.7e-03 |
| Uchl3 | 175.3 | 93.8 | -0.9 | 1.2e-05 |
| Cnnm2 | 35.8 | 174.9 | 2.3 | 8.6e-08 |
| Cxxc5 | 174.3 | 111.0 | -0.6 | 6.5e-03 |
| Gpr63 | 112.5 | 169.5 | 0.6 | 8.5e-04 |
| Llgl1 | 167.6 | 116.8 | -0.5 | 2.2e-03 |
| Cnrip1 | 165.6 | 117.7 | -0.5 | 9.5e-03 |
| Thumpd2 | 165.3 | 97.2 | -0.8 | 6.5e-03 |
| Slco3a1 | 22.6 | 161.2 | 2.9 | 7.5e-14 |
| Ggcx | 158.3 | 119.9 | -0.4 | 5.9e-03 |
| Chaf1b | 156.6 | 102.7 | -0.6 | 4.6e-03 |
| Ubtd1 | 106.7 | 156.4 | 0.6 | 3.2e-04 |
| Gdf11 | 153.5 | 76.7 | -1.0 | 8.1e-08 |
| Arrdc4 | 72.4 | 151.4 | 1.1 | 1.5e-07 |
| Hdac6 | 151.1 | 95.1 | -0.7 | 7.0e-04 |
| Gm21190 | 149.5 | 69.4 | -1.1 | 3.4e-04 |
| Glis2 | 147.9 | 77.5 | -0.9 | 3.1e-03 |
| Cbarp | 63.3 | 145.2 | 1.2 | 6.1e-07 |
| Pfkfb4 | 141.3 | 94.8 | -0.6 | 3.7e-03 |
| Mrm1 | 101.1 | 139.0 | 0.5 | 3.4e-03 |
| Rarb | 55.0 | 138.5 | 1.4 | 3.9e-07 |
| Pold1 | 138.0 | 89.2 | -0.6 | 4.8e-03 |
| Ears2 | 52.0 | 137.3 | 1.4 | 1.3e-05 |
| Atp5mk | 89.4 | 135.5 | 0.6 | 5.8e-03 |
| Klhl25 | 95.5 | 134.7 | 0.5 | 1.6e-03 |
| Abhd11 | 130.9 | 92.0 | -0.5 | 8.8e-03 |
| Rab11fip5 | 77.5 | 125.4 | 0.7 | 2.1e-03 |
| Sestd1 | 87.4 | 124.6 | 0.5 | 3.2e-03 |
| Gpr85 | 80.9 | 120.8 | 0.6 | 4.0e-03 |
| Bcar3 | 116.3 | 52.9 | -1.1 | 4.6e-03 |
| Osbpl1a | 113.6 | 80.7 | -0.5 | 7.1e-03 |

Continued on next page

Table S2 – continued from previous page

| Gene Name | CONTROL | ADO (2h) | log2FC | padj |
|---|---|---|---|---|
| Nckap5l | 111.4 | 73.9 | -0.6 | 6.2e-03 |
| Rgl3 | 107.8 | 70.9 | -0.6 | 5.9e-03 |
| Sh2b2 | 39.5 | 104.0 | 1.4 | 2.8e-05 |
| Hes1 | 98.8 | 36.3 | -1.5 | 2.3e-03 |
| Qsox2 | 62.1 | 98.1 | 0.6 | 4.1e-03 |
| Pcgf2 | 98.0 | 53.8 | -0.9 | 9.4e-05 |
| Ano8 | 96.4 | 67.5 | -0.5 | 6.4e-03 |
| Igsf5 | 16.5 | 95.0 | 2.5 | 3.4e-12 |
| Focad | 93.7 | 62.2 | -0.6 | 3.9e-03 |
| Tgfa | 29.9 | 93.3 | 1.7 | 1.4e-05 |
| Pde10a | 23.2 | 86.7 | 1.9 | 2.2e-04 |
| Pdlim1 | 32.4 | 83.7 | 1.4 | 8.5e-04 |
| Snn | 82.3 | 51.8 | -0.7 | 8.6e-03 |
| Kcnd1 | 80.5 | 47.6 | -0.8 | 8.9e-03 |
| Klra2 | 36.5 | 76.8 | 1.1 | 1.2e-04 |
| Polg2 | 75.5 | 50.4 | -0.6 | 9.7e-03 |
| Slc35f3 | 74.8 | 44.0 | -0.7 | 7.1e-03 |
| Ccsap | 54.2 | 73.1 | 0.5 | 5.3e-03 |
| P2ry10 | 29.0 | 71.6 | 1.3 | 2.6e-03 |
| Mcm10 | 70.9 | 43.1 | -0.7 | 2.3e-03 |
| Zfp248 | 70.1 | 40.0 | -0.8 | 4.2e-03 |
| Snx7 | 66.8 | 47.2 | -0.5 | 2.4e-03 |
| Slc46a1 | 23.5 | 64.9 | 1.5 | 7.3e-05 |
| C1qtnf12 | 25.6 | 61.6 | 1.2 | 3.7e-04 |
| Gm20696 | 56.2 | 24.6 | -1.2 | 1.6e-04 |
| Msi1 | 54.5 | 29.1 | -0.9 | 4.8e-03 |
| Ncaph | 53.9 | 30.0 | -0.8 | 5.5e-03 |
| Lrrc36 | 53.7 | 18.4 | -1.6 | 3.8e-03 |
| Hoxb4 | 51.9 | 32.9 | -0.6 | 8.0e-03 |
| Sdhaf4 | 49.9 | 29.1 | -0.8 | 9.1e-03 |
| Atp8b1 | 48.8 | 20.8 | -1.2 | 3.3e-04 |
| Draxin | 20.8 | 46.6 | 1.2 | 1.1e-05 |
| Fancd2 | 45.9 | 27.8 | -0.7 | 6.4e-03 |
| Twist2 | 45.3 | 15.8 | -1.6 | 1.5e-03 |
| Dennd5b | 43.5 | 19.1 | -1.2 | 7.3e-03 |
| Rnf227 | 42.9 | 24.0 | -0.9 | 1.7e-04 |
| Lpar2 | 26.4 | 42.6 | 0.7 | 6.1e-03 |
| Hid1 | 13.3 | 42.3 | 1.7 | 9.6e-04 |
| Whrn | 13.0 | 41.9 | 1.7 | 7.4e-04 |
| Frmd5 | 16.4 | 41.0 | 1.3 | 2.7e-03 |
| Sap30 | 18.9 | 36.9 | 0.9 | 4.3e-03 |

Continued on next page

Table S2 – continued from previous page

| Gene Name | CONTROL | ADO (2h) | log2FC | padj |
|---|---|---|---|---|
| Hfe | 14.7 | 36.6 | 1.3 | 1.0e-03 |
| Micall2 | 12.4 | 35.0 | 1.5 | 1.0e-03 |
| Mettl15 | 34.5 | 20.8 | -0.7 | 8.8e-03 |
| Nphp4 | 9.0 | 34.4 | 2.0 | 6.5e-05 |
| Grhpr | 32.6 | 15.4 | -1.1 | 5.8e-03 |
| Slc25a35 | 29.1 | 12.1 | -1.2 | 5.5e-03 |
| Tifa | 27.7 | 13.3 | -1.2 | 5.2e-03 |
| Lrrc2 | 11.1 | 26.9 | 1.3 | 7.3e-03 |
| Samd11 | 7.7 | 26.1 | 1.7 | 3.0e-03 |
| Rwdd2a | 25.8 | 8.3 | -1.7 | 1.4e-04 |
| Or2y1f | 22.5 | 4.2 | -2.4 | 1.8e-04 |
| Dagla | 4.5 | 21.3 | 2.3 | 1.5e-04 |
| Gng4 | 6.1 | 20.2 | 1.8 | 3.1e-03 |
| Rln3 | 1.3 | 17.3 | 4.0 | 1.4e-06 |
| Fndc9 | 16.8 | 6.0 | -1.2 | 3.6e-03 |
| Bmp2 | 16.4 | 5.5 | -1.5 | 2.6e-03 |
| Il7 | 5.4 | 16.1 | 1.4 | 3.0e-03 |
| Apol9b | 16.1 | 4.7 | -1.9 | 6.7e-03 |
| Itgb6 | 15.8 | 5.6 | -1.7 | 4.5e-03 |
| Me3 | 4.1 | 15.4 | 1.8 | 1.6e-03 |
| Tmem202 | 5.6 | 14.1 | 1.4 | 5.4e-04 |
| Trim58 | 12.1 | 2.4 | -2.3 | 1.6e-05 |
| Kcns3 | 10.7 | 2.3 | -2.2 | 1.6e-04 |

**Table S3:** Summary of top six most abundant protein classes among upregulated ADO-specific genes.

| Protein Classes | Associated Genes |
|---|---|
| **metabolite interconversion enzyme (PC00262)** | |
| acyltransferase(PC00042) | Elovl5 Cpt1a |
| aldolase(PC00044) | Aldoa |
| carbohydrate kinase(PC00065) | Pfkp |
| deacetylase(PC00087) | Nceh1 |
| dehydrogenase(PC00092) | Impdh1 Sdhb Ldha |
| galactosidase(PC00104) | Galc |
| guanylate cyclase(PC00114) | Gucd1 |
| kinase(PC00137) | Pik3cb Dgke Pi4kb |
| lipase(PC00143) | Dagla Golt1b |
| nucleotide phosphatase(PC00173) | Entpd7 |
| oxidase(PC00175) | Qsox2 |
| oxidoreductase(PC00176) | Me3 |
| phosphodiesterase(PC00185) | Prune1 Pde10a |
| phospholipase(PC00186) | Pla2g4a |
| transferase(PC00220) | Hs6st1 |
| **protein-binding activity modulator (PC00095)** | |
| G-protein(PC00020) | Arf2 |
| GTPase-activating protein(PC00257) | Arhgap25 Sipa1l1 |
| guanyl-nucleotide exchange factor(PC00113) | Trio Cyth1 |
| heterotrimeric G-protein(PC00117) | Gng4 |
| kinase activator(PC00138) | Ccnc |
| kinase modulator(PC00140) | Prkar1a |
| protease inhibitor(PC00191) | Serpinb6 Lxn |
| small GTPase(PC00208) | Rab27b Rhoq Rap2b Rab11fip5 Rab4a |
| **transporter (PC00227)** | |
| ATP synthase(PC00002) | Atp5f1d Atp5mk |
| ion channel(PC00133) | Cnnm2 |
| primary active transporter(PC00068) | Mfsd12 |
| transporter(PC00227) | Kpna1 Slco3a1 Slc16a3 Pitpnc1 Slc20a2 Tnpo1 |
| voltage-gated ion channel(PC00241) | Cbarp Kcnn4 Vdac3 |
| **cytoskeletal protein (PC00085)** | |
| actin binding motor protein(PC00040) | Myo5a Tpm4 |
| actin or actin-binding cytoskeletal protein(PC00041) | Pdlim1 Arpc3 |
| cytoskeletal protein(PC00085) | Tmem202 Whrn |
| microtubule binding motor protein(PC00156) | Dctn4 |
| microtubule or microtubule-binding cytoskeletal protein(PC00157) | Wdr90 |

Continued on next page

Table S3 – continued from previous page

| Protein Classes | Associated Genes |
|---|---|
| non-motor actin binding protein(PC00165) | Frmd5 Scin Coro1c |
| **transmembrane signal receptor (PC00197)** | |
| G-protein coupled receptor(PC00021) | Grm5 Gpr63 Lpar2 Mrgprx2 Gpr85 P2ry10 |
| transmembrane signal receptor(PC00197) | Tlr4 Csf2rb2 Fzd6 Fzd7 Csf2rb |
| **scaffold_adaptor protein (PC00226)** | |
| scaffold/adaptor protein(PC00226) | Rassf3 Ywhaz Sntb2 Micall2 Klhl23 Mpp7 Mfhas1 Sh2b2 Plin2 Arrb1 |

**Table S4:** Summary of top two most abundant protein classes among downregulated ADO-specific genes.

| Protein Classes | Associated Genes |
|---|---|
| **gene-specific transcriptional regulator (PC00264)** | |
| C2H2 zinc finger transcription factor(PC00248) | Glis2 Zfhx2 Zeb1 Zfp619 Zfp248 Ikzf2 |
| DNA-binding transcription factor(PC00218) | Gata1 Nfix Gata2 |
| HMG box transcription factor(PC00024) | Cic |
| basic helix-loop-helix transcription factor(PC00055) | Hes1 |
| homeodomain transcription factor(PC00119) | Hoxb4 |
| transcription cofactor(PC00217) | Crtc1 Arid4b |
| winged helix/forkhead transcription factor(PC00246) | Rfx5 Rfx7 |
| **protein modifying enzyme (PC00260)** | |
| cysteine protease(PC00081) | Pgpep1 Uchl3 Usp20 |
| non-receptor serine/threonine protein kinase(PC00167) | Ksr1 Taok3 Map4k2 Stk39 Prkcd Mink1 |
| protein modifying enzyme(PC00260) | P4ha1 |
| serine protease(PC00203) | Mbtps1 |
| ubiquitin-protein ligase(PC00234) | Pias2 Trim58 Itch Rnf227 |

**Table S5:** Summary of genes involved in inflammatory mediator biosynthesis and their receptors in mast cells

| Category | Associated Gene (Ligand) | Receptor(s) |
| --- | --- | --- |
| ***Lipid Mediator Synthesis*** | | |
| Precursor Liberation | Pla2g4a, Pla2g6, Dagla, Daglb, Napepl | |
| Cyclooxygenase | Ptgs1, Ptgs2, Hpgds, Ptgds, Ptges, Ptges2, Ptges3, Tbxas1 | |
| Lipoxygenase | Alox5, Alox5ap, Ltc4s, Lta4h, Alox12, Alox15 | |
| Platelet-Activating Factor | Lpcat1, Lpcat2 | |
| Cytochrome P450 | Cyp2j5, Cyp2j6 | |
| ***Cytokines, Chemokines, & Growth Factor*** | | |
| Chemokine Superfamily | Ccl2 | Ccr2 |
| | Ccl3 | Ccr1, Ccr5 |
| | Ccl4 | Ccr5 |
| | Ccl5 | Ccr1, Ccr3, Ccr5 |
| | Ccl7 | Ccr1, Ccr2, Ccr3 |
| | Ccl8 | Ccr2 |
| | Ccl11 | Ccr3 |
| | Ccl12 | Ccr2 |
| | Ccl24 | Ccr3 |
| | Cxcl1 | Cxcr2 |
| | Cxcl2 | Cxcr2 |
| | Cxcl3 | Cxcr2 |
| | Cxcl5 | Cxcr2 |
| | Cxcl9 | Cxcr3 |
| | Cxcl10 | Cxcr3 |
| | Cxcl11 | Cxcr3 |
| | Cxcl12 | Cxcr4 |
| | Xcl1 | Xcr1 |
| | Cx3cl1 | Cx3cr1 |
| Interleukin (IL) Family | Il1a, Il1b, Il1f10 | Il1r1/Il1rap |
| | Il2, Il4, Il7, Il9, Il15 | Il2rg (common $\gamma$-chain) |
| | Il3 | Il3ra/Csf2rb/Csf2rb2 |
| | Il4, Il13 | Il4ra/Il13ra1/Il2rg |
| | Il5 | Il5ra/Csf2rb/Csf2rb2 |
| | Il6 | Il6r/Il6st |
| | Il9 | Il9r/Il2rg |
| | Il10 | Il10ra/Il10rb |

*Continued on next page*

Table S5 – *Continued from previous page*

| Category | Associated Gene (Ligand) | Receptor(s) |
|---|---|---|
| | Il12a, Il12b | Il12rb1/Il12rb2 |
| | Il17a, Il17f | Il17ra/Il17rc |
| | Il18 | Il18r1/Il18rap |
| | Il19, Il20, Il24 | Il20ra/Il20rb |
| | Il22 | Il22ra1/Il10rb |
| | Il23a, Il12b | Il23r/Il12rb1 |
| | Il27 | Il27ra/Il6st |
| | Il31 | Osmr/Il31ra |
| | Il33 | Il1rl1 |
| | Il34 | Csf1r |
| TNF Superfamily | Tnf | Tnfrsf1a, Tnfrsf1b |
| | Lta | Tnfrsf1a |
| | Tnfsf4 | Tnfrsf4 |
| | Tnfsf9 | Tnfrsf9 |
| | Tnfsf10 | Tnfrsf10b |
| | Fasl | Fas |
| | Cd40lg | Cd40 |
| Interferon (IFN) Family | Ifna (all subtypes), Ifnb1 | Ifnar1/Ifnar2 (Type I) |
| | Ifng | Ifngr1/Ifngr2 (Type II) |
| | Ifnl2, Ifnl3 | Ifnlr1/Il10rb (Type III) |
| TGF-$\beta$ Superfamily | Tgfb1, Tgfb2, Tgfb3 | Tgfbr1/Tgfbr2 |
| | Bmp2, Bmp4, Bmp7 | Bmpr1a/Bmpr1b/Bmpr2/Acvr2a/Acvr2b |
| | Inhba, Inhbb | Acvr1b/Acvr2a/Acvr2b |
| | Tgfa | Egfr |
| CSFs | Csf1 | Csf1r |
| | Csf2 | Csf2ra/Csf2rb/Csf2rb2 |
| | Csf3 | Csf3r |
| Wnt Family | Wnt1, Wnt2, Wnt2b, Wnt3, Wnt3a, Wnt4, Wnt5a, Wnt5b, Wnt6, Wnt7a, Wnt7b, Wnt8a, Wnt8b, Wnt9a, Wnt9b, Wnt10a, Wnt10b, Wnt11, Wnt16 | Fzd1-10, Lrp5, Lrp6 |
| Other Growth Factors | Areg | Egfr |
| | Vegfa | Kdr, Flt1 |
| | Vegfb | Flt1 |
| | Pdgfa, Pdgfb | Pdgfra/Pdgfrb |
| | Edn1 | Ednra |
| | Flt3l | Flt3 |
| | Kitl | Kit |

*Continued on next page*

Table S5 – *Continued from previous page*

| **Category** | **Associated Gene (Ligand)** | **Receptor(s)** |
|---|---|---|
| | Lif | Lifr |
| | Osm | Osmr |
| | Thpo | Mpl |
| | Tslp | Crlf2/Il7r |
| Other Secreted Proteins | Hmgb1 | Ager, Tlr2, Tlr4 |
| | Mif | Cd74 |
